# Examining Litter Specific Variability in Mice and its Impact on Neurodevelopmental Studies

**DOI:** 10.1101/2022.09.09.506402

**Authors:** Vanessa Valiquette, Elisa Guma, Lani Cupo, Daniel Gallino, Chloe Anastassiadis, Emily Snook, Gabriel A. Devenyi, M. Mallar Chakravarty

**Affiliations:** Integrated Program in Neuroscience, McGill University, Montreal, QC, Canada; Department of Biology and Biomedical Engineering, McGill University, Montreal, QC, Canada; Department of Psychiatry, McGill University, Montreal, QC, Canada; Computional Brain Anatomy Laboratory, Cerebral Imaging Centre, Douglas Mental Health University Institute, Montreal, QC, Canada; Institute of Medical Science & Collaborative Program in Neuroscience, University of Toronto, Toronto, ON, Canada; Department of Medicine, University of Toronto, Toronto, ON, Canada

**Author notes:** Author contributions: EG, DG, CA, and ES data acquisition; EG and GB image processing and MAGeT pipeline; VV data analysis; VV paper redaction; EG, LC, DG, GB and MMC paper edits. Funding: Canadian Institutes for Health Research (CIHR), Healthy Brains for Healthy Lives (HBHL) and Fonds de Recherche du Quebec en Sante (FRQS).

## Abstract

Our current understanding of litter variability in neurodevelopmental studies using mouse may limit translation of neuroscientific findings. Higher variance of measures across litters than within, often termed intra-litter likeness, may be attributable to pre- and postnatal environment. This study aimed to assess the litter-effect within behavioral assessments (2 timepoints), and anatomy using T1-weighted magnetic resonance images (4 timepoints) across 72 brain region volumes (36 C57bl/6J inbred mice; 7 litters: 19F/17M). Between-litter comparisons of brain and behavioral measures and their associations were evaluated using univariate and multivariate techniques. A power analysis using simulation methods was then performed modeling neurodevelopment and evaluating trade-offs between number-of-litters, mice-per-litter, and sample size. Our results show litter-specific developmental effects, from the adolescent period to adulthood for brain structure volumes and behaviors, and their associations in adulthood. Our power simulation analysis results suggest increasing the number-of-litters in experimental design to achieve the smallest total sample size for detecting different rates of change in specific brain regions. Our results also demonstrate how litter-specific effects may influence development and that increasing the litters to the total sample size ratio should be strongly considered when designing neurodevelopmental studies.

## 1. Introduction

Although inbred mouse strains may share genetics, there is increasing acknowledgement that individual mice within a strain exhibit significant variance in brain structure, function and behavior. Factors such as maternal care, litter size, intrauterine enviroment, sex (Crews et al., 2009; Golub & Sobin, 2020; Meyer et al., 2009; Qiu et al., 2018) and postnatal environment may impact critical periods of early development (Golub & Sobin, 2020; McCarty, 2017; Meyer et al., 2009) where different neurobiological processes (e.g. neurogenesis, synaptogenesis, corticogenesis, etc) affect gray and white matter development (Keshavan & Paus, 2015; Lebel & Beaulieu, 2011; Paus et al., 2008). Environmental exposures both in utero and early life can impact these processes and the resulting neuroanatomical developmental trajectories (Gogtay & Thompson, 2010; Guma, do Couto Bordignon, et al., 2021). Microstructural changes occurring in response to these exposures may cumulatively lead to neurodevelopmental morphological variation (e.g. brain structure volume) that can be captured by magnetic resonance imaging (MRI) (Lerch et al., 2017; Shaw et al., 2008).

Studies of the impact of environmental exposures on neurodevelopment often use rodent models given the tight control experimenters have on genetics and the housing environment. Despite this experimental control, the pup’s litter of origin could be a key component, as factors such as litter size and sex-ratio may increase inter-litter variance, thereby dwarfing potential biological effects (Festing, 2006; Golub & Sobin, 2020; Jiménez & Zylka, 2021; Lazic & Essioux, 2013; Zorrilla, 1997). From a statistical standpoint, the independence of measures is not satisfied for pups from the same litter, causing some experimenters to use the litter itself as the sample of interest (Lazic et al., 2018; Lazic & Essioux, 2013). In this scenario, it is common to average the measures by litter or randomly select one pup per litter. While these are potentially desirable design choices, they are an inefficient use of resources. Using a linear mixed effect model (LMER) which allows for a hierarchical statistical design, is potentially one of the best suggested ways to account for inter-litter variance (Flood et al., 2012; Lazic & Essioux, 2013).

There have been previous attempts to examine this potential litter-effect (Festing, 2006; Hughes, 1979; Wainwright, 1998; Zorrilla, 1997) and this interest has recently re-emerged (Golub & Sobin, 2020; Lazic & Essioux, 2013; Vorhees & Williams, 2021) resulting in guidelines aimed to normalize study design and methodological reporting as means of improving reproducibility and replicability (Vorhees & Williams, 2021) (ARRIVE [(Percie du Sert et al., 2020)], maternal immune activation (MIA) guidelines [(Kentner et al., 2019)], and Festing & Altman’s guidelines [(Festing & Altman, 2002)]). These studies are further motivated by recent observations that offspring exposed to MIA (modeling the exposure to maternal infection during pregnancy, a known risk factor for neurodevelopmental disorders) show litter-dependent variance in resilience and susceptibility to the risk factor exposure (Mueller et al., 2021). However, our group also demonstrated that the gestational timing of MIA-exposure could further modulate neurodevelopmental outcomes (Guma, do Couto Bordignon, et al., 2021; Guma et al., 2022; Guma, Snook, et al., 2021). At an even more basic level, litter-dependent modulation of body and brain weight have been reported using linear and linear mixed-effect models; these effects are further affected by sex (Golub & Sobin, 2020; Jiménez & Zylka, 2021). Thus, there is a critical need to identify how different experimental design choices are indeed litter-dependent and their putative impact on measures of brain anatomy or behavioral measures (Golub & Sobin, 2020).

Our limited understanding and inconsistent handling of litter likely acts as an experimental confound (Jiménez & Zylka, 2021; Lazic & Essioux, 2013; Leenaars et al., 2019), and has been linked to increased rates of false positives (Williams et al., 2017). This study aimed to examine putative effects of litter on neuroanatomy (derived using magnetic resonance imaging) and behavior aimed at capturing changes relevant to neurodevelopmental disorders. Specifically, we seek to study the following questions: 1) Can we observe brain and behavior differences within or between litters in normative mouse development? 2) Is there a specific pattern of regional brain volume that can explain litter-specific variance in mouse offspring? 3) Do any of the brain region patterns covary with patterns of behaviors? and 4) When modeling neurodevelopment, how does the number-of-litters and the litter size impact the ability to detect within-subject and between-group regional volume changes?

## 2. Methods

This project builds upon previously acquired and analyzed data from our group studying the impact of prenatal MIA-exposure on brain and behavioral development in mice (Guma, do Couto Bordignon, et al., 2021). 1) Our first step was to evaluate normative development via the observation of a control group. We evaluated the litter variance and median differences for each brain structure volume and for each behavior using the Brown-Forsythe-Levene’s (BFL) and Kruskal-Wallis (KW) tests. 2) Subsequently, we detected patterns of brain regions that covary together using Principal Components Analysis (PCA). 3) The possible relationship between behavioral measures and PCA-derived patterns of brain anatomy were further analyzed using Partial Least Squares Correlation (PLSC). 4) Using Linear Mixed Effects Models (LMER), we examined models of normative neurodevelopment. Models goodness-of-fit were examined with and without controlling for the litter. 5) A power-analysis using a simulation-based technique was performed according to the best selected model for normative neurodevelopment and group differences (treatment-like vs control). The power simulation aimed to assess the best trade-off between the number-of-litters to the number of mice per litter (litter-size) required to observe significant differences in longitudinal regional brain volume.

### 2.1 Animals

We selected data acquired from mice derived from dams injected with saline (SAL; gestational day (GD) 9 or 17, 0.9% sterile NaCl solution) (Fig. 1) as previously reported (Guma, do Couto Bordignon, et al., 2021) (42 total C57bl/6J inbred mice; 9 litters: 20F / 22M; 2 litters only one sex). This former project was approved by McGill University’s Animal Care Committee under the guidelines of the Canadian Council on Animal Care. Mice were maintained under a 12-hour light cycle (8am-8pm), with food and water access ad libitum. From the total sample, 2 litters were dismissed due to the insufficient number of offspring and data per timepoint (number mice/litter ≤ 3), resulting in a total of 36 mice selected (17M/19F ; 7 litters; 2 litters only one sex; Fig. 2; Supplementary Table S1). Selected mice were weaned and sexed at postnatal day (PND) 21 and housed with same-sex siblings for a total of 2-4 mice per cage. Subsequently, mice were scanned at PND 21, 38, 60 and 90 and underwent a series of behavioral tests following the PND 38 and 90 imaging timepoints (Fig. 1). Our earliest timepoint, PND 21, will be refered as childhood, PND 38 as adolescence, PND 60 as early adulthood and PND 90 as adulthood. There is still debate regarding development period timeline, and in this case, PND 60 could reflect a transitional period between adolescence and adulthood (Hammelrath et al., 2016; Semple et al., 2013).

**Fig. 1 :**
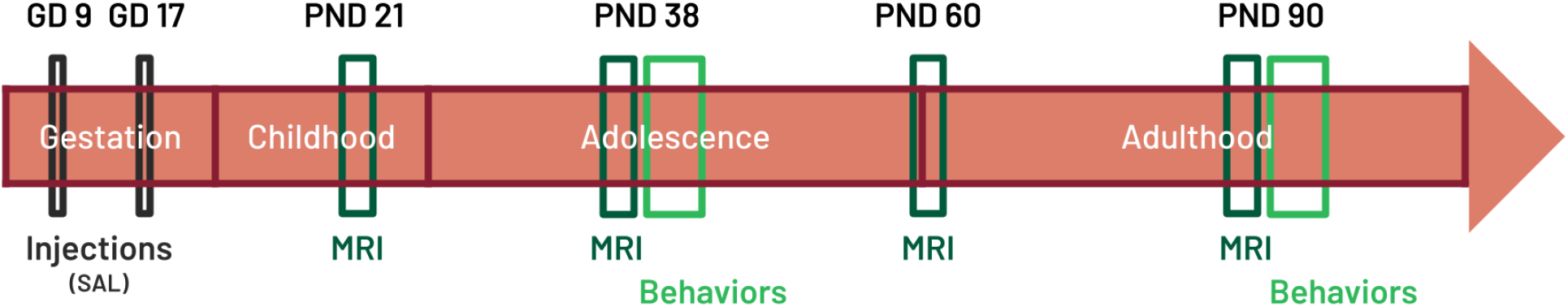
Data acquisition timeline from the dataset acquired in a previous MIA study from our group. The data selected consists of control mice whose mothers were injected with SAL at GD 9 or 17. Mice were scanned at PND 21, 38, 60 and 90 and underwent a series of behavioral tests following PND 38 and 90 scans including the marble burying, prepulse inhibition and open field tests (Guma, do Couto Bordignon, et al., 2021).

**Fig. 2 :**
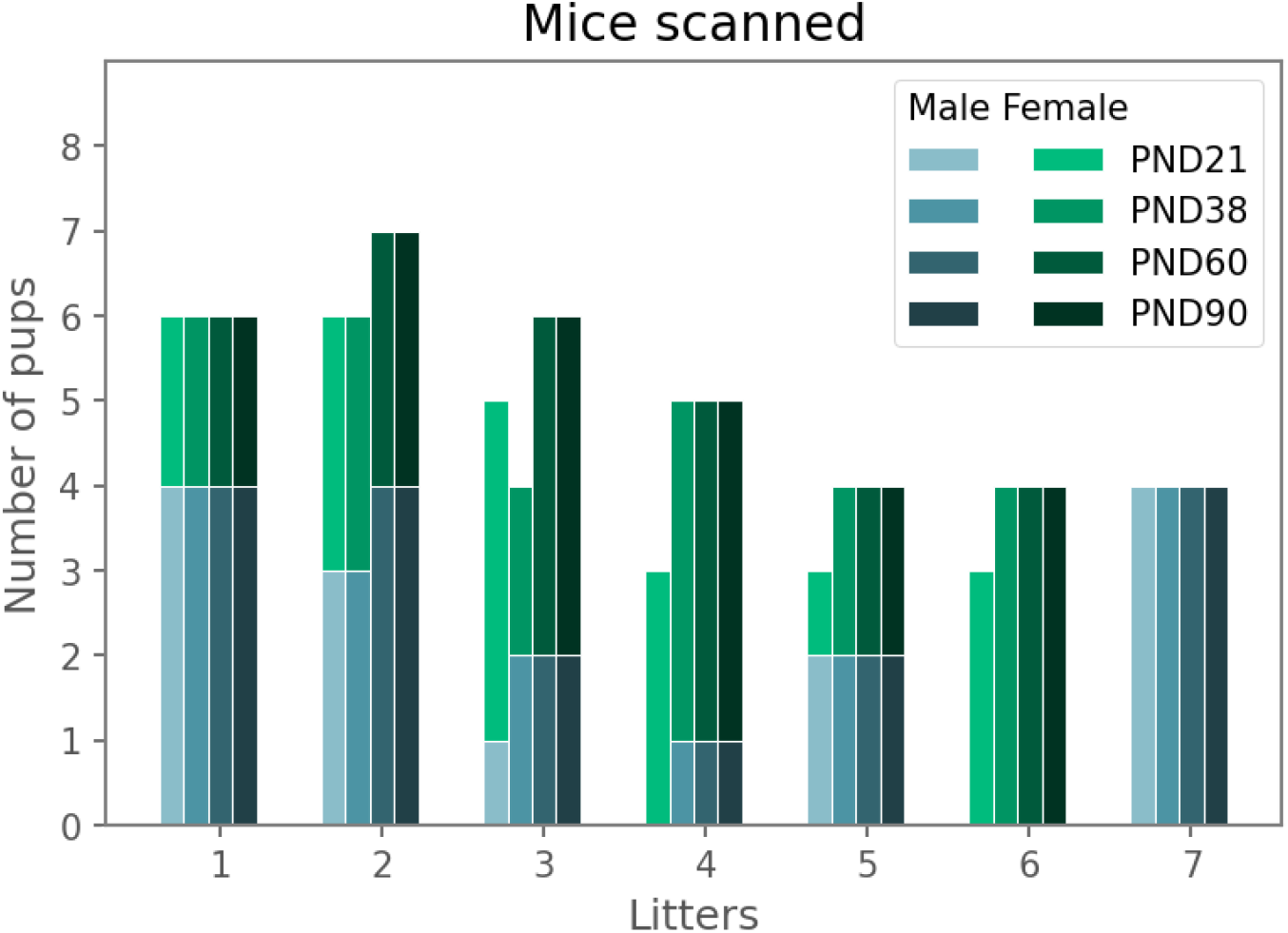
Distribution of scanned mice by sex, litter and timepoint.

### 2.2 Behavioral testing

Tests were previously selected and assessed based on the impact of MIA, a potent risk factor for neurodevelopmental disorders, on offspring behaviors at PND 38 and 90. Tests were performed 1-2 days following scans and with a 1-2 days rest period between tests to minimize stress (Guma, do Couto Bordignon, et al., 2021). These tests included: the marble burying task (MBT) to assess stereotypical/repetitive behaviors, the prepulse inhibition test (PPI) to evaluate filtering of unnecessary information and sensorimotor gating, and the open field test (OFT) to assess locomotion, anxiety-like and exploratory behaviors. The measures selected from each test are : the number of marbles 100% buried, and 75% buried from the MBT; the maximum and average startle response to a 120 decibels (dB) stimulus for the first and last 5 trials, and following a prepulse of 0, 3, 6, 9, 12, 15 dB during the main set from the PPI; the duration, frequency and distance relative to edges, corners and center zone, and overall total distance from the OFT. Note that for the PPI, the average measures reflects the average startle amplitude and the maximum measures reports the maximum startle amplitude recorded for a specific prepulse stimulus. Specifics regarding these procedures are described in supplemental (Supplementary Method S1) and in greater details elsewhere (Guma, do Couto Bordignon, et al., 2021).

### 2.3 Magnetic resonance imaging

Anatomical in vivo brain images (Structural, 100um^3^ voxels, T1-weighted images, MnCl_2_ injection [50 mg/kg] 24 hours prior for contrast enhancement, matrix size of 180 x 160 x 90, 14.5 min, 2 averages; 5% isoflurane for induction, 1.5% for maintenance) were previously acquired at PND 21, 38, 60 and 90 with a 7T Bruker Biospec 70/30 MRI scanner for small animals (Guma, do Couto Bordignon, et al., 2021) (Fig. 2).

Once MRI images were acquired, several preprocessing steps were performed and described in greater details elsewhere (see (Guma, do Couto Bordignon, et al., 2021). Briefly, DICOM images were converted to MINC files, denoised (Coupe et al., 2008), corrected with the N4ITK algorithm for the inhomogeneity of the bias field (Tustison et al., 2010) and registered with a rigid-body alignment (Friedel et al., 2014). Visual quality control (QC) was performed to reject scans presenting artifacts or anomalies (e.g. motion, signal drop off, hydrocephalus), as described in (Guma, do Couto Bordignon, et al., 2021).

The Pydpiper multiple automatically generated templates (MAGeT) brain segmentation pipeline (https://github.com/CoBrALab/MAGeTbrain; (Chakravarty et al., 2013; Friedel et al., 2014)), a multi-atlas registration-based segmentation tool, was used to automatically segment mice neuroanatomy from preprocessed T1-weighted images. Segmentations were based on the DSURQE atlas (Dorr-Steadman-Ullmann-Richards-Qiu-Egan, https://wiki.mouseimaging.ca/display/MICePub/Mouse+Brain+Atlases) (Dorr et al., 2008; Qiu et al., 2018; Richards et al., 2011; Steadman et al., 2014; Ullmann et al., 2013). A total of 356 region volumes were initially extracted from the segmentation labels. Labels were hierarchically merged to form a superset of 72 structure volumes (including 36 region labels for both hemispheres [VERM_R and VERM_L labels overlap partially on vermal subregions and differ on paravermal regions] ; https://github.com/CoBrALab/documentation/wiki/DSURQE-atlas-hierarchical-downsample, Supplementary Table S2) to ease interpretation in our exploratory analyses.

### 2.4 Litter variation in normative development analysis

To focus on normative litter-variation over the course of development we used the saline-exposed control group. Brain and behavioral measures were standardized by timepoint using the StandardScaler() function from scikit-learn package (v0.24.1) in Python (v3.8.6). Once these measures were standardized, we first evaluated the differences in variance and median of regional brain volume and behavioral measures between litters (univariate analysis). Second, PCA-derived brain patterns were estimated to determine if litter was related to components of variance (multivariate analysis). Finally, we examined how PLSC-derived patterns of interlinked covariance attributable to brain and behavioral variables may be related to litter origin (multivariate analysis) (Fig. 3).

**Fig. 3 :**
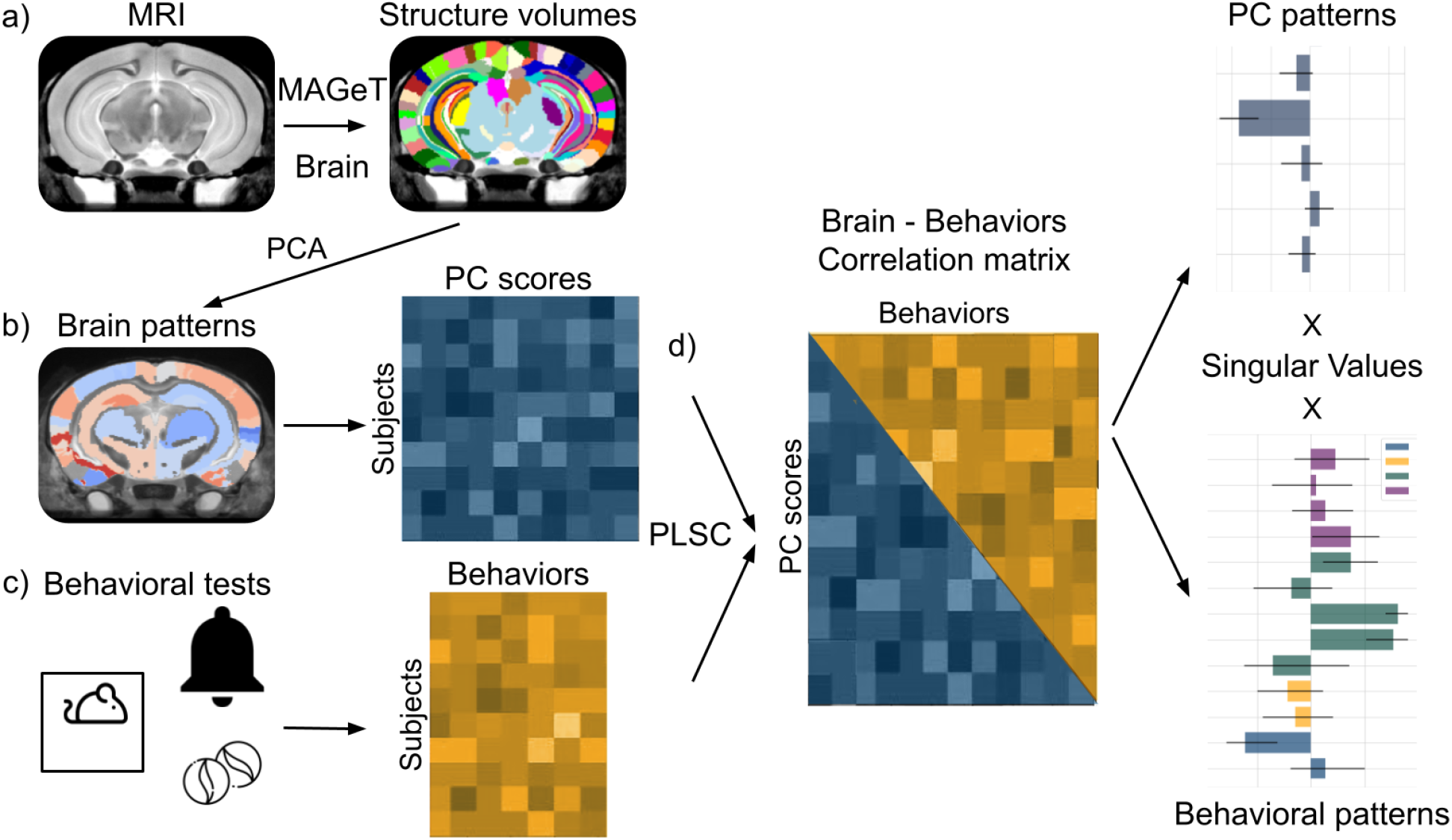
Multivariate analysis workflow performed at every timepoint (PND 21, 38, 60, 90). a) Brain volumes were initially extracted following MAGeT brain segmentation (356 labels) of MRI images and grouped to create 72 structure volumes. b) Volumes were z-scored and used in PCA to find patterns of regional brain volumes and collected in a PC scores matrix. c) Z-scored behavioral assessments, and descriptive data (sex and litter size) were collected in a behavioral matrix. d) The PC scores were then used as a derived brain measure matrix in PLSC against the behavioral matrix. This step conferred covariation patterns of PC scores and behaviors, resulting in specific PC scores patterns and behavioral patterns for each explanatory latent variable. (Structure volumes image adapted from : https://wiki.mouseimaging.ca/display/MICePub/Mouse+Brain+Atlases)

#### 2.4.1 Univariate analysis : Litter distribution

According to the Shapiro-Wilk test results (shapiro() from SciPy package (v1.7.3) in Python) (Supplementary Table S3) and our small sample size, nonparametric tests of variability and median examinations were selected. The BFL equality of variance test (stats.levene() from SciPy package in Python) (Brown & Forsythe, 1974), was used to evaluate if homogeneity of variance was present between litters for every brain and behavioral measure at each timepoint. A significant result (p < 0.05) indicated that one or more groups had different variances. The KW test (stats.kruskal() from SciPy package in Python) was then used to compare median differences in brain volume and behavioral measures between litters. A significant result (p < 0.05) indicated that one or more litters had different medians. Since these analyses were applied over multiple variables, false discovery rate (FDR) correction (stats.multitest.fdrcorrection() from statsmodel package (v0.12.1) in Python) (Benjamini & Hochberg, 1995) was used to detect the proportion of false positives (type 1 error) at a 5% threshold.

#### 2.4.2 Multivariate analysis: Principal component analysis

Principal Components Analysis (PCA), was used to assess if litter membership explained any part of data-driven components of variance observed in specific brain structures’ patterns. PCA transforms a dataset with high-dimensionality, potentially correlated variables, into a matrix of orthogonal Principal Components (PC) that explains most of the variance. Brain volumes (72 regions), extracted from the MAGeT brain segmentation pipeline, that covary together were estimated seperately for every available timepoint. Subsequently, PC scores litter-specific variance and medians were evaluated by using BFL and KW tests.

The syndRomics package (v0.1.0) in R (v4.1.3) (https://github.com/ucsf-ferguson-lab/syndRomics) (Torres-Espín et al., 2021) was used to perform PCA on z-scored brain structure volumes. Thereafter, permutation testing (n=1000) was performed on PCs and loadings followed by a 5% threshold FDR correction (Benjamini & Hochberg, 1995) across all brain areas. Stability of results was assessed with bootstrap resampling (n=1000) to estimate the confidence interval of the PCs loadings.

#### 2.4.3 Multivariate analysis: Partial least squares correlation

A Partial Least Squares correlation (behavioral) analysis (PLSC) (Abdi & Williams, 2013; Krishnan et al., 2011) was used to evaluate components modeling the covariance between brain anatomy and behaviors. This correlational approach aims to find maximal covariance between 2 matrices, without any directionality (prediction) from one matrix to the other by estimating Latent Variables (LV), where LVs are ordered by the amount of covariance explained. For every timepoint were behavioral and brain measures were acquired, we retained the top PCs that cumulatively explained 95% of the total variance (PND38: 22PCs, PND 90: 24PCs) in an aim to maximize the number of components included but by discarding PC’s explaining a trivial portion of variance. These PC scores were included in the PLSC to assess covariance of brain patterns (Total of 22 or 24 variables) related to behaviors. Behavioral measures included: MBT, the number of marbles buried, either fully (100%) or partially (75%) were included (2 variables); PPI maximum measures for 0, 3, 6, 9, 12, 15 dB prepulse (6 variables); OFT measures for time spent, frequency of entries, and distance traveled in the edges, the corners, and the center zone of the arena, and total distance traveled in the whole arena (10 variables) (see section 2.2). Descriptive data such as sex and litter size (2 variables) were added to the behavioral matrix to evaluate their associations. Selecting PCs as input variables for this analysis gave the model initial insight in potential brain structure patterns relevant at a particular timepoint and provided a means to reduce the dimensionality of the dataset, an important consideration in multivariate analyses. Previous behavioral measures results (Guma, do Couto Bordignon, et al., 2021) and our current BFL, and KW results (see section 3.1) guided the selection of behaviors to further include in the PLSC. The pyls package (v0.1.6) in Python was used for PLSC (https://github.com/rmarkello/pyls). Permutation testing (n=5000) was performed to evaluate significance of LVs and bootstrapping (n=5000) to evaluate reliability of the LVs via confidence intervals.

### 2.5 Power simulation of mice neurodevelopment

For this section, we sought to evaluate appropriate sample size selection for longitudinal studies while considering the impact of litter variability on the ability to observe statistically significant results for a given effect size. We first performed the power simulation based on the total collection of regional brain volumes normative development (within-subject change) lmer models with varied effect sizes. Then, from the control group, a treatment group was simulated by duplicating the control subjects and increasing their brain structure volumes by 1%. This allowed the maintenance of the properties of the distribution within sample by only simulating a between group difference. A second power simulation was performed based on group differences and through a range of simulated effect sizes. For this purpose, the control group only will be referred to as the control data and the control and treatment simulated groups will be referred to as the treatment data.

#### 2.5.1 Linear mixed effects modeling volumetric trajectories according to litter grouping

Linear mixed effects models are ideal for capturing changes in longitudinal study designs (Bernal-Rusiel et al., 2013) and are commonly used in longitudinal neuroimaging studies (Chakravarty et al., 2015; Guma, do Couto Bordignon, et al., 2021; Qiu et al., 2018). The integration of grouping effects (random effects) allows a certain control of the unexplained residual variance that could be originating from variables modulating main results. It is especially important for hierarchical data, where statistical independence of observation is violated (Brauer & Curtin, 2018; Schielzeth & Nakagawa, 2013; Wainwright et al., 2007). The lmer function from the lme4 package (v1.1-28) in R (Bates et al., 2015) was used to model brain structure volumes changes through time (PND 21, 38, 60, 90) with different factors for the control data:

model 1: Brain region ∼ sex + age + (1|mouse)
model 2: Brain region ∼ sex * age + (1|mouse)
model 3: Brain region ∼ sex + age + (1|mouse) + (1|litter)
model 4: Brain region ∼ sex * age + (1|mouse) + (1|litter) and to model group differences through time for the treatment data :
model 5: Brain region ∼ treatment group * age + sex + (1|mouse) + (1|litter)

To further examine the extent to which a model with or without the litter as a random effect best suited our data, the Akaike Information Criterion (AIC) (Bozdogan, 1987) was evaluated by using the aictab function from the AICcmodavg package (v2.3.1) in R (https://cran.r-project.org/web/packages/AICcmodavg/AICcmodavg.pdf) only for the control data.

#### 2.5.2 Power simulation of normative and treatment-like neurodevelopment

A population simulation-based power analysis, inspired by (Lerch et al., 2012), was performed to evaluate power for varying combinations of litter-size, number-of-litters, and sample size. This was implemented using the R2power function within the mixedpower package (v0.1.0) in R (Brysbaert & Stevens, 2018; Green & MacLeod, 2016; Kumle et al., 2021) to perform a 2 level (litter-size, number-of-litters) power simulation. The power simulation for regional brain volume changes in time, modeled from controls according to model 3 with all variables in a numeric form, was based on the results from every brain region LMER for 3 timepoints (PND 38, 60, 90). PND 21 data entries were omitted due to a significant number of mice with fewer acquired measures compared to later timepoints. For a more streamline simulation, only mice with data entries at every timepoints were used. This new dataset was further extended by randomly duplicating subjects (Fig. 4 a-b), until the maximum sample size, litter-size and number-of-litters we wanted to simulate was reached. The maximum litter-size was fixed at 6 mice to mimic the average litter-size of C57bl/6 dams in preclinical settings. A single power simulation round was completed for a simulated (n=1000) number-of-litters [3-14] and fixed number of mice per litter (litter-size) (e.g. 2), and ran subsequently for every iteration of litter-size [2-6], for a fixed age effect size. Simulation rounds were completed for simulated age effect sizes of 0.10, 0.15, 0.20, or 0.25 for the control data and, from the treatment data to model group differences based on model 5 (see section 2.5.1) and [4-30] number-of-litters (in increments of 2) and [2-6] mice per litter (litter-size), for a 0.35, 0.40, 0.45 treatment group*age effect size. Power estimates for every litter-size by number-of-litters combination, were plotted in a heatmap for every brain region (72 regions) and for every simulated effect size. From those heatmaps, 2 types of best combination were extracted:

1. Smallest Sample size (SS). From every heatmap power estimates, the best litter-size by number-of-litters trade-off achieving the smallest sample size (SS) and reaching > 80% power was selected. This optimizes study design from a waste-reduction perspective of overall sample size.
2. Maximized number of mice by litter (MM). From every heatmap power estimates, the first trade-off that reached > 80% power and maximized the number of mice per litter (MM) was selected. This method mimics the standard practice for selecting sample size in a laboratory setting.

**Fig. 4 :**
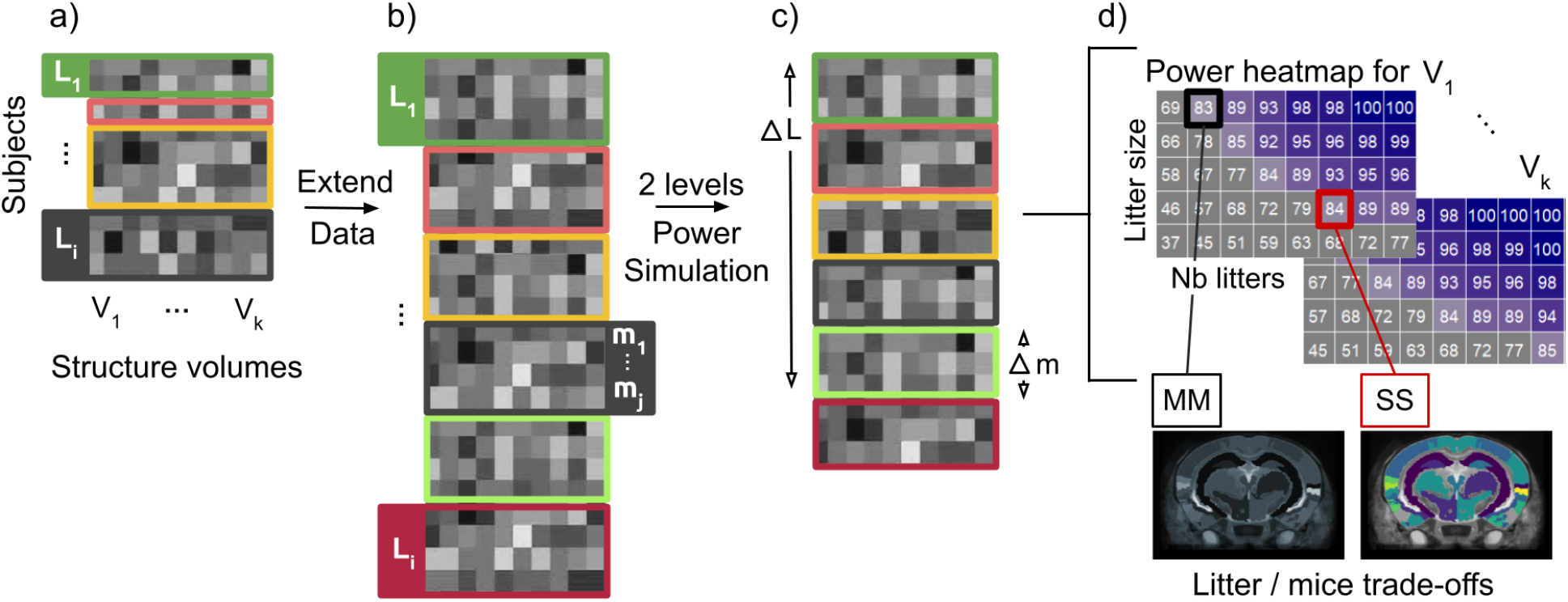
Power simulation workflow. a) The initial input was the brain structures volume (V_k_) matrix containing measures from subjects (mice) nested within litters (L) with corresponding number-of-mice (litter-size, m). b) A new extended dataset was simulated to reach a fixed number of litters (L_i_) and a fixed number of mice per litter (m_j_). c) The power simulation was performed, with this new extended matrix, and simulated along 2 levels, number(nb)-of-litters (ΔL) and number-of-mice-per-litter (Δm). d) For every brain region, the power estimates were gathered in a power heatmap. The best trade-offs (MM and SS) were selected for every brain structure’s heatmap and projected onto a summary brain map.

The SS results selected for each brain region were then mapped onto a standard mouse brain average for every simulated effect size run (Fig. 4).

## 3. Results

### 3.1 Normative development

#### 3.1.1 Litter variation

We observed mainly trend-level differences in litter-specific variance, based on the BFL test, for both brain and behavior measures (Fig. 5, Fig. 6, Supplementary Table S4). For brain structure volumetry, differences in litter variance were observed at PND 38 for the left primary somatosensory area, the right piriform-amygdalar area, and only significant after FDR correction for the left thalamus (Fig. 5). At PND 60, differences were observed, but not maintained after FDR correction, for the right auditory and the left agranular insular/orbital areas. No litter-specific differences were observed across measures at PND 21 or PND 90. Despite the modest litter-specific variance observed within a handful of brain regions, a tendency for greater variance is highlighted during the adolescence and early adulthood period.

**Fig. 5 :**
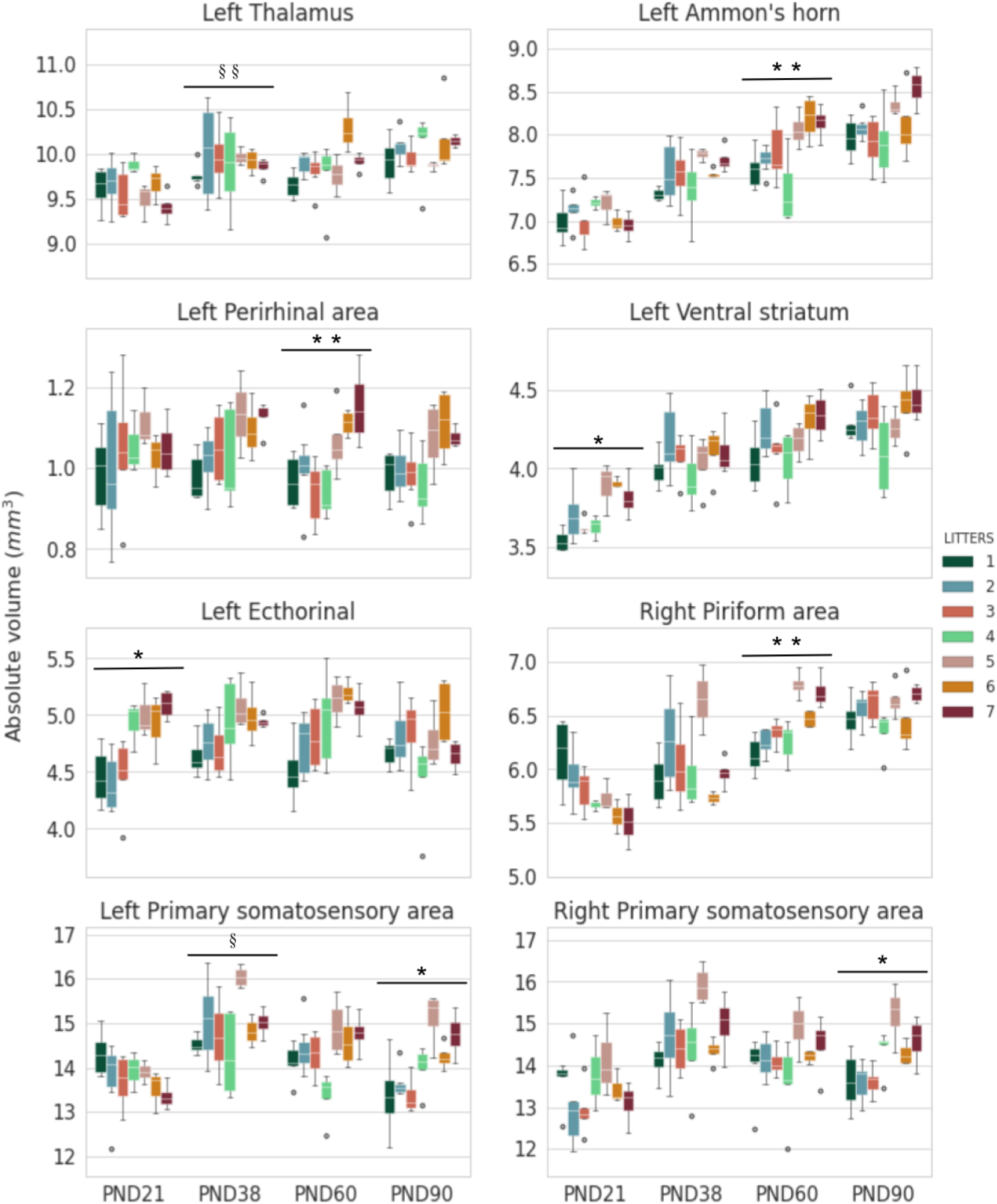
Distribution of brain region volumes by litters. Regions shown here were selected to demonstrate the most representative BFL and KW results. Presented here are observed variance heterogeneity (BFL) between litters at PND 38 for the left thalamus and left primary somatosensory area. Litter median differences (KW), for the left ectorhinal area at PND 21, for the left ammon’s horn, the left perirhinal area, the right piriform area at PND 60, and for the primary somatosensory bilateral areas at PND 90. (BFL : sig. = §, after FDR=§§ ; KW: sig. = *, after FDR=**)

**Fig. 6 :**
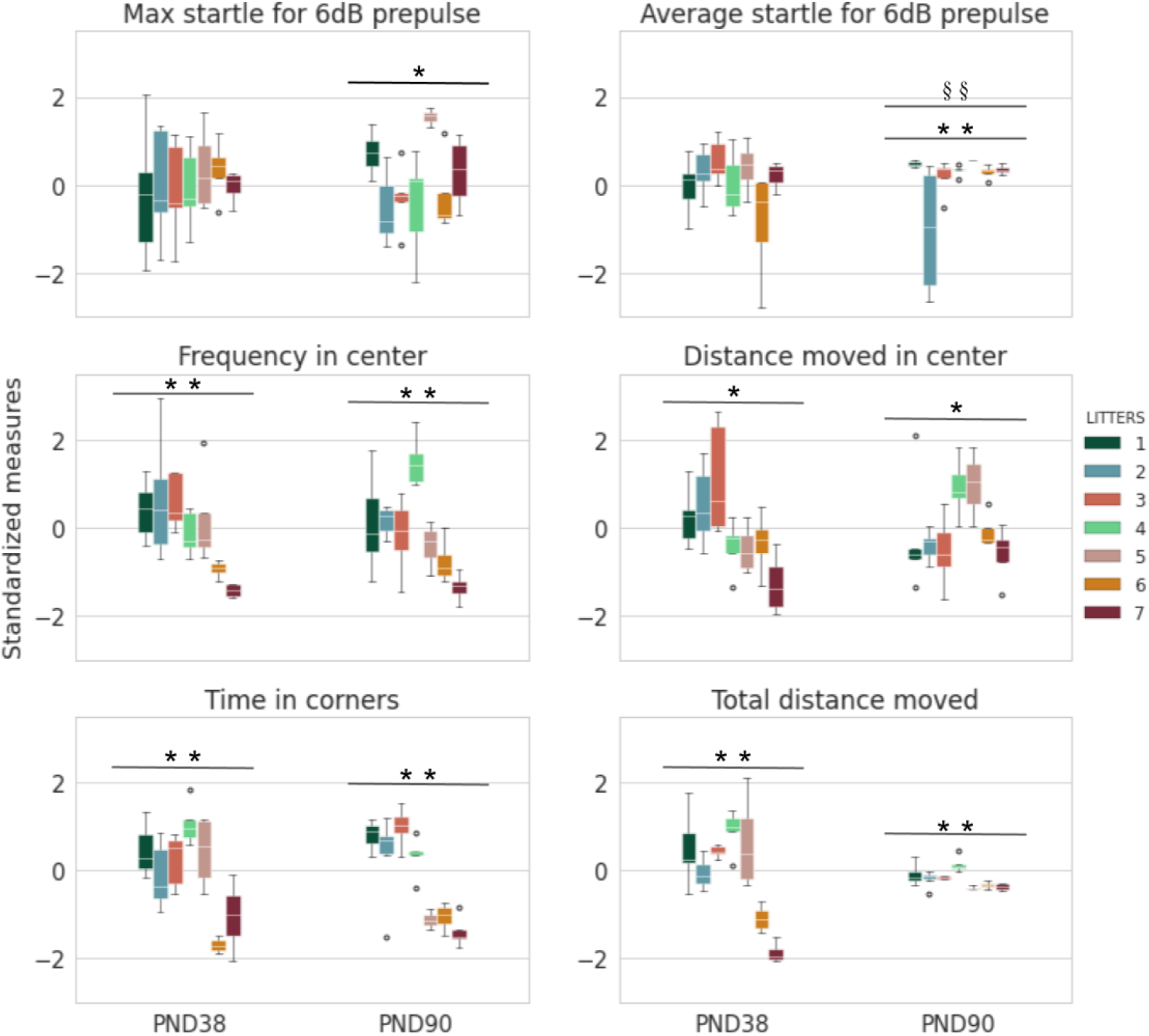
Standardized behavioral measure distributions by litters. We can observe variance heterogeneity (BFL) for the average startle response for trials with a 6 dB prepulse, and median differences (KW) between litters at PND 38, for the frequency and distance moved in the center, time in the corners and total distance, and at PND 90 for averaged and maximum startle response for a 6 dB prepulse, frequency and distance moved in the center, time in corners and total distance. (BFL : sig. = §, after FDR=§§ ; KW: sig. = *, after FDR=**)

For behavioral measurements, at PND 38, greater litter variation was observed in OFT measurements for the cumulative duration in the center of the arena, and for the PPI test average startle response of trials without prepulse measured at the end of the test. At PND 90, we observed that all the PPI average trial data from the different prepulse intensities (3, 6, 9, 12, 15 dB) had significant differences in litter variance (Supplementary Table S4). After further investigation, BFL results seemed to be driven by one litter (litter 2) at this specific timepoint, which had a much greater variance compared to other litters (PPI average 6 dB, Fig. 6). This specific litter had a distinct grouping between males and females for results pertaining to the PPI test compared to other litters that had more homogeneous measures within them. PPI average measures only, based on those results, were not carried to the following steps of the analysis. No differences were observed for all MBT measurements. Taken together, some litter-specific variance was observed for behavioral measurements, specifically the OFT, during the adolescent period.

We observed litter-specific median differences, based on KW test, for both brain and behavior measures. For brain structure volumes, subthreshold differences were observed at PND 21, for the left ecthorinal cortex and ventral striatum. At PND 60, significant differences after FDR correction were observed in the left and right dorsal striatum, the left and right hippocampus ammon’s horn, the left hemispheric cerebellar region, the right piriform area, and the left perirhinal area. Finally, at PND 90, subthreshold effects were observed in the left and right primary somatosensory areas and the left cortical subplate (claustrum, amygdala) (Fig. 5, Supplementary Table S5). No significant results were observed at PND 38. Most litter-specific brain volumes differences are observed in early adulthood.

Differences were observed in behavioral measures litter-specific medians at PND 38 for the distance and frequency moved across the entire arena for the corners, edges, and center, as well as total, and cumulative duration in the center and edges for the OFT. For the PPI test, average trials measures for the 9 dB startle response also showed differences in litter medians. At PND 90 the litter-related differences in OFT behaviors persisted. Similarly, for PPI average startle amplitude we observed differences in litter medians for measures across prepulse intensities [6, 9, 12, 15 dB], maximum startle amplitude without prepulse at the end and average at the beginning, and maximum startle at 6 dB (Fig. 6, Supplementary Table S5). MBT measurements did not show litter differences. Differences were observed for all timepoints where behavioral measures were acquired, both during adolescence or adulthood, especially for the OFT measurements of anxious-like and exploratory behaviors.

#### 3.1.2 Brain patterns

Our PCA results demonstrated one PC at PND 38 and three PCs at each of the other timepoints surviving permutation testing and FDR correction. At PND 21, the three significant PCs explained 24.5% (PC1, p_perm_=0.001, q=0.002), 11.7% (PC2, p_perm_=0.001, q=0.002) and 9.4% (PC3, p_perm_=0.001, q=0.002) of the variance (Supplementary Fig S1). PC1 included significant smaller regions (negative loadings), across the cerebellum, hippocampal region (dentate gyrus), hypothalamus, thalamus, and somatosensory areas, amongst others (Fig. 7, Supplementary Fig S2). PC2 (KW, H=21.622, p=0.001, q=0.022) expressed a sparser pattern of smaller regions (negative loadings) for the left temporal association areas, the ventral striatum, the ectorhinal cortex, and the auditory areas, and the frontal pole bilaterally, and bigger regions (positive loadings) for the left cortical and piriform-amygdalar areas and the piriform areas bilaterally (Fig. 7, Supplementary Fig S2). The PC2 scores showed litter-specific distributions scattered along either side of the null axis in line with the subsequent KW test revealing differences between litter distribution medians (Fig. 8). PC3’s most significant negative components (smaller regions) were the right ventral striatal and the ectorhinal cortex, and positive (bigger regions) were the left perirhinal area and the cortical subplate (claustrum, amygdala) (Fig. 7, Supplementary Fig S2).

**Fig. 7 :**
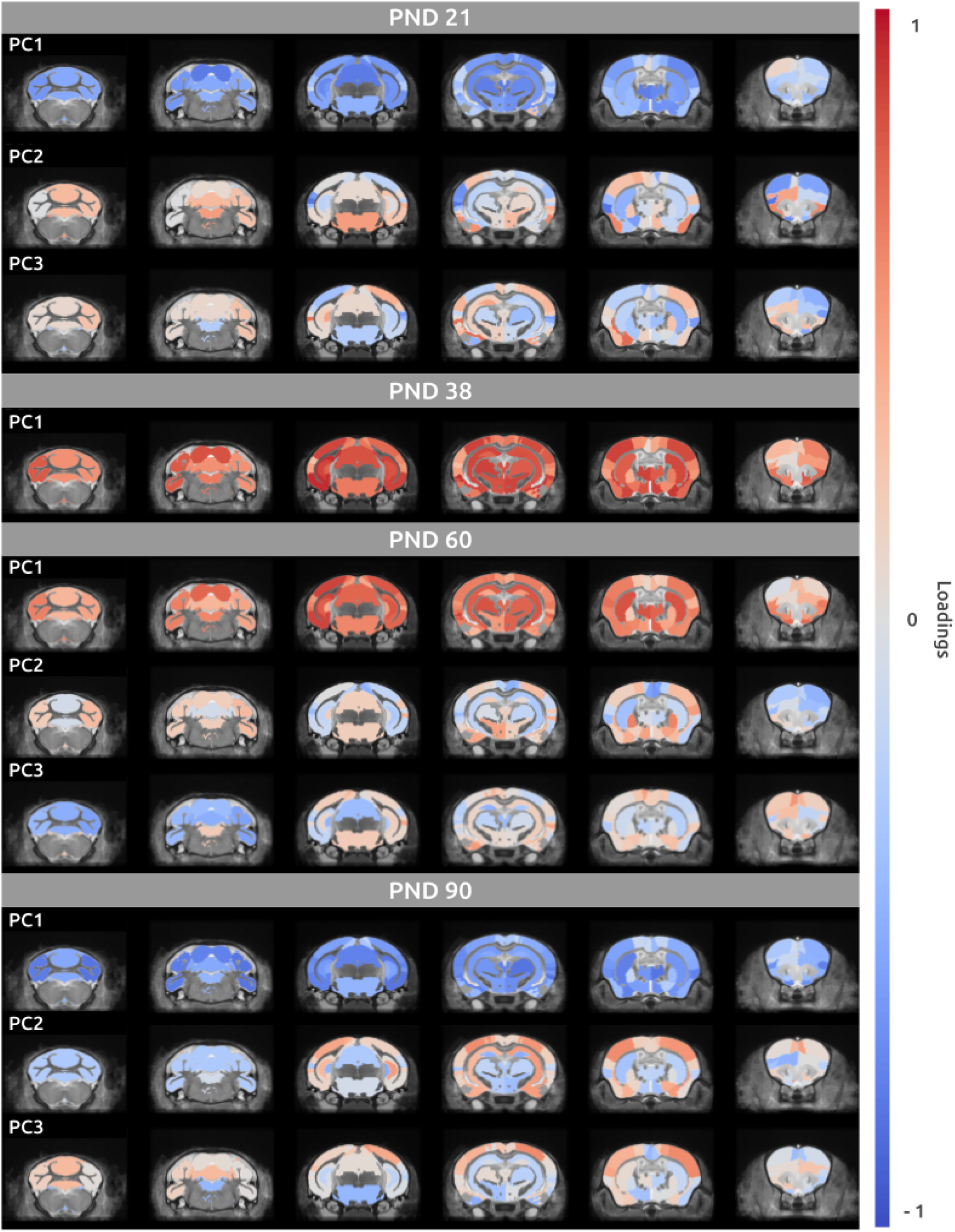
PCA loadings maps. For every significant principal component the associated loadings were overlaid on average mouse brain template.

**Fig. 8 :**
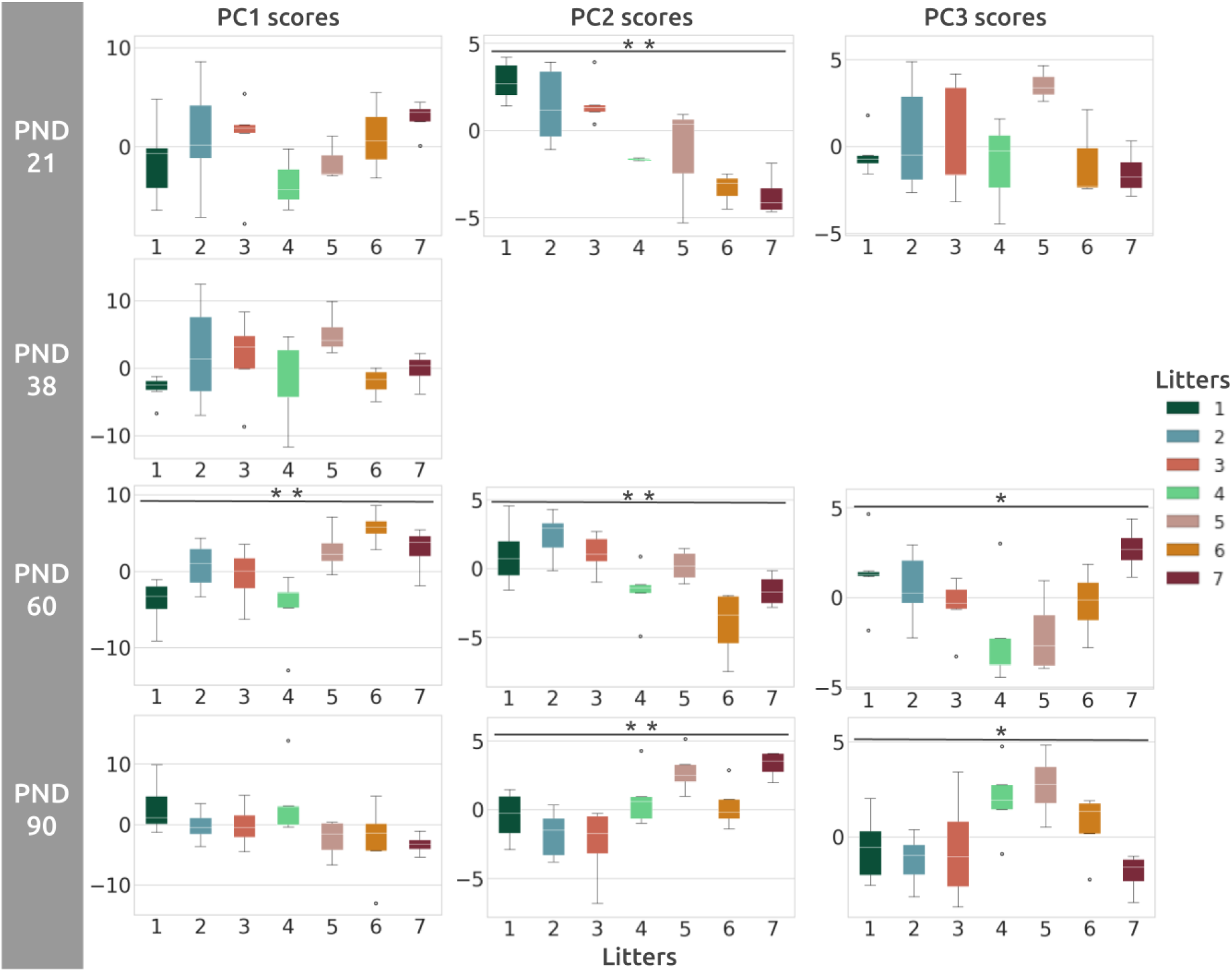
PCA scores distributions by litters for every individual timepoint. For every significant principal component, PC scores distributions are shown per litter and their significance relative to BFL and KW testing. (KW: sig. = *, after FDR=**)

At PND 38, PC1 explained 41.1% (p_perm_=0.001, q=0.005) of the overall variance and was the only significant component covering almost the entire brain anatomy (Supplementary Fig S1). The retrohippocampal regions, the cortical subplate (claustrum, amygdala), the thalamus, the striatal and the hippocampal regions contributed positively (bigger regions) to this component (Fig. 7, Supplementary Fig S2). According to PC1 scores distributions by litter, there was no observable evident litter grouping associated with PC1 (Fig. 8) and potentially coding for overall brain size at this age (Narvacan et al., 2017; Shaw et al., 2008).

At PND 60, PC1, PC2, and PC3 each explained 29.3% (p_perm_=0.001, q=0.002), 9.5% (p_perm_=0.001, q=0.002), and 8.4% (p_perm_=0.001, q=0.002), and their respectives scores demonstrated clear differences in litter-specific contributions (Fig. 8, Supplementary Fig. S1). PC1 (KW, H=20.271, p=0.002, q=0.026) significant larger regions (positively contributing regions) were mainly associated with the visual regions, the thalamus, the striatum, the retrohippocampal and the hippocampal (Ammon’s horn) regions, and the hindbrain (Fig. 7, Fig. 8, Supplementary Fig. S2). Depicting similar regions also observed in PC1 at PND 38, and most likely showing greater overall brain structure growth spurt in adolescence (Narvacan et al., 2017; Shaw et al., 2008). For PC2 (KW, H=21.491, p=0.001, q=0.023), the right piriform-amygdalar area, the secondary motor area, and the anterior cingulate cortex bilaterally were negatively (smaller regions) contributing to this component and inversely, positively loading (bigger regions) were the right supplemental somatosensory area, and the bilateral dorsal regions of the pallidum (Fig. 7, Fig. 8, Supplementary Fig S2). For PC3 (KW, H=14.442, p=0.025), one predominant region and its subregions loaded negatively (smaller), such as bilaterally the cerebellar nuclei, the vermal and the hemispheric areas of the cerebellum and a handful of regions loading positively (bigger regions) barely achieved significance (Fig. 7, Fig. 8, Supplementary Fig S2).

At PND 90, PC1, PC2, and PC3 explained 29.2% (p_perm_=0.001, q=0.002), 9.6% (p_perm_=0.001, q=0.002), and 7.2% (p_perm_=0.001, q=0.002) of the variance, respectively (Supplementary Fig S1). PC1 was driven by negative loadings (smaller regions) for the thalamus, the striatum, the retrohippocampal region, the temporal association areas, the hemispheric regions of the cerebellum, the midbrain and the right hindbrain (Fig. 7, Fig. 8, Supplementary Fig S2). PC2 scores showed litter differences (KW, H=19.911, p=0.003, q=0.045) and this PC was associated with strong negative loadings (smaller regions) for the cerebellar nuclei, the dentate gyrus, and the left agranular insular/orbital areas, and positive (bigger regions) for the primary somatosensory and the motor areas, the piriform-amygdalar and the cortical subplate (claustrum, amygdala) (Fig. 7, Fig. 8, Supplementary Fig S2). PC3 scores were also driven by litter differences (KW, H=15.169, p=0.019), and this component had really specific association with regions pertaining to the striatum-like amygdalar nuclei and the bilateral cortical amygdalar area, loading negatively (smaller regions), and the primary, the supplemental somatosensory and the posterior parietal association areas contributing positively (bigger regions) (Fig. 7, Fig. 8, Supplementary Fig S2).

Overall, PC1s from every timepoint depicted a widespread brain pattern highlighting overall smaller regions at PND 21 and 90 and bigger ones at PND 38 and 60. At PND 21, the left ventral striatum and the ectorhinal regions exhibited litter-specific variance (KW, Fig. 5, see section 3.1.1, Supplementary Table S5) and were part of PC2 that also exhibited litter-specific scores differences (KW, Fig. 8). Similar associations were seen at PND 60, where the left hemispheric cerebellar region (KW, see section 3.1.1, Supplementary Table S5) was associated with greater volume in PC1 and smaller volume in PC3 (Fig. 7, Supplementary Fig. S2), and were all litter-dependant (KW, Fig. 8). Finally, at PND 90, only the left and the right primary somatosensory areas (KW, Fig. 5, see section 3.1.1, Supplementary Table S5) were positively associated with PC2 and 3 (Fig. 7, Supplementary Fig. S2), also reflecting litter-specific variances (KW, Fig. 8).

#### 3.1.3 Brain and behaviors patterns

At PND 38, LV2 was statistically significant (p_perm_ = 0.034) and explained 17.8% of the covariance (Supplementary Fig. S3). This latent variable did not covary significantly with any of the previous litter-specific PCs (Supplementary Fig. S4).

At PND 90, LV1, 2, and 4 were statistically significant (p_perm_= [0.014, 0.005, 0.087]) and explained respectively 32.3%, 16.9% and 9.5% of the covariance (Supplementary Fig. S3). LV1 and LV4 did not show clear covariance between behaviors and litter-specific PCs (see section 3.1.2, Supplementary Fig. S5). LV2 (KW, H=18.869, p=0.004) (Fig. 9b) showed specific positive covariances with PC2 and 3 (loadings: 0.406, 0.681) (Fig. 9a), both previously revealing litter-dependent relationships (see section 3.1.2, Fig. 8). More specifically, PC2 and PC3 represented smaller cerebellar nuclei, allocortex (dentate gyrus, left agranular insular/orbital areas, right striatum-like amygdalar nuclei) and cortical amygdalar regions and bigger primary, supplemental somatosensory and motor regions, and piriform-amygdalar and posterior parietal association areas (Fig. 8, Supplementary Fig S2). These patterns were positively correlated with specific measures corresponding to the center zone (OFT) and negatively to the amount of time in corners (OFT) and the litter size (Fig. 9a). Taken together, mice exhibiting greater PC2 and/or 3 brain patterns (smaller cerebellar nuclei, striatum-like amydgalar nuclei and dentate gyrus, and bigger cortical subplate, somatosensory areas and piriform-amygdalar regions) would also show greater exploratory and less anxious-like behaviors, and be from a smaller litter (Fig. 9a).

**Fig. 9 :**
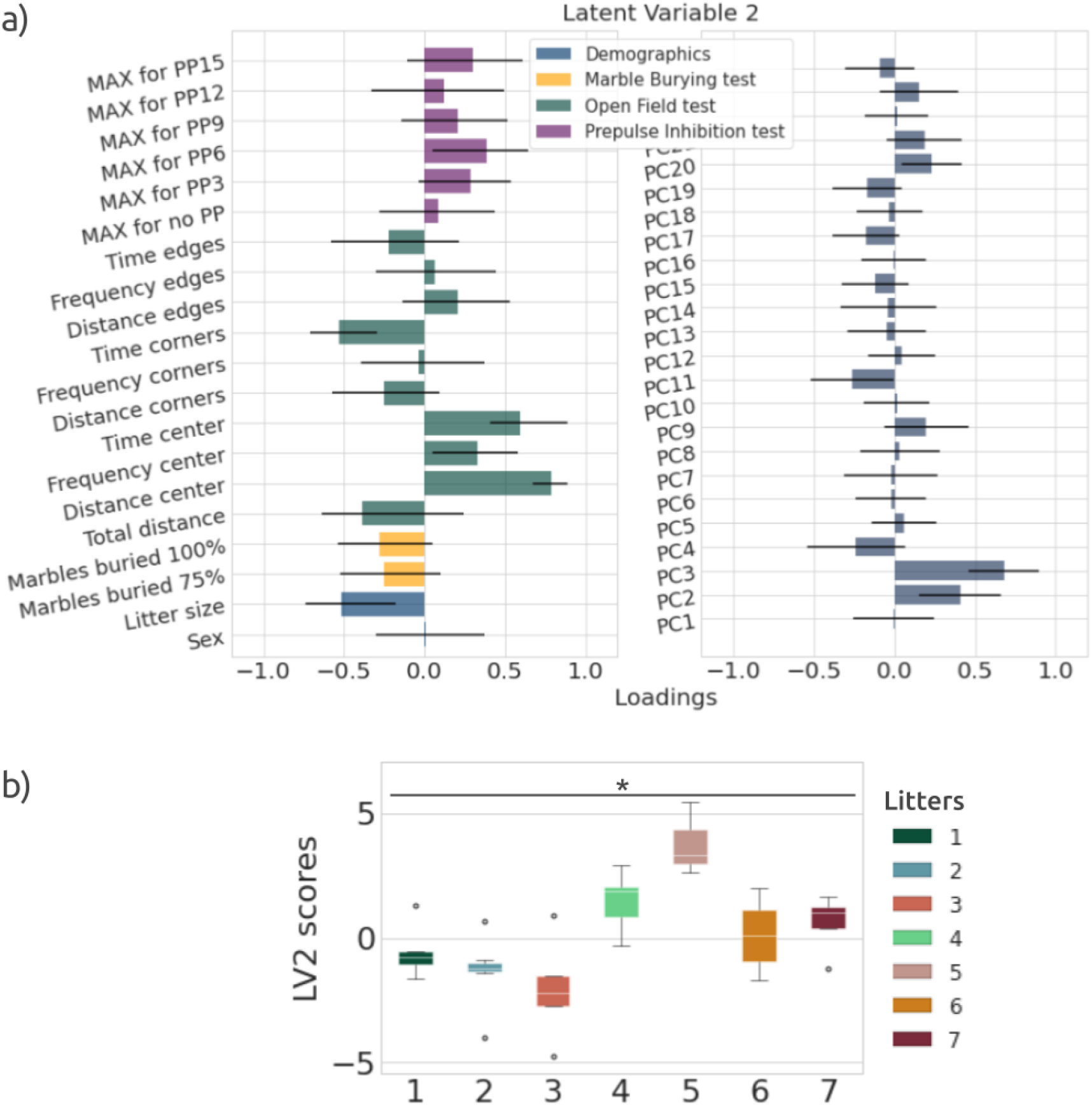
Behaviors and PCs contributions to LV2 at PND 90 and its scores distribution by litters. a) LV2 is positively associated with PC2-3 brain patterns (loadings: 0.406, 0.681), and measures of OFT relative to the center zone. The litter size and the time in corners (OFT) were negatively associated. b) LV2 scores distributions per litter. LV2 score medians are significantly different between litters. (KW : sig.= *, after FDR=**)

To summarize, these results suggest litter-dependent variations for individual brain volumetric and behavioral measures over the developmental period studied here, mainly observable starting in adolescence (univariate analysis, KW and BFL, Fig. 5 and Fig. 6, Supplementary Table S4 and Table S5). They are also showing litter-dependent integrative measurements presented in form of brain patterns, for every timepoint except adolescence (multivariate analysis, PCA, Fig. 8), presenting different region associations, and their covariation with behaviors, only in adulthood (multivariate analysis, PLSC, Fig. 9, Supplementary Fig. S4 and Fig. S5).

### 3.2 Power simulation of normative and treatment-like mice development

#### 3.2.1 Linear mixed effects model

Four mixed effect models were examined for each brain region to model normative development. Model 1 integrated a fixed effect of sex and age and a random effect of mouse and model 2 added an interaction between sex and age. Model 3 and model 4 were the same as model 1 and 2, respectively, with the addition of a litter random effect. When comparing every model’s AIC for each region, 7 regions benefited from model 3 and 8 from model 4 out of 72 regions (based on lower AIC values) (Table 1). Furthermore, adding the litter as a random effect, in cases where the region benefited from this model (lower AIC values) (Table 1), a modulation of the sex effect was also observed (Table 2). From these highlighted regions, the ventral striatum, the piriform, the ectorhinal, the frontal pole, the auditory, and the temporal association areas were also represented in previous analysis results (BFL, KW, PCA, PLSC, see section 3.1) from childhood, and the somatosensory regions, cortical amygdalar, and temporal association areas represented in adulthood (Table 1). These specific brain regions associated with lower AIC results for model 3 and model 4 overlapped with regions present in brain patterns (PCs) showing a litter-effect at PND 21 (PC2) and at PND 90 (PC3) (Table 1).

**Table 1 :** AIC results of brain growth model comparisons w/ and w/o litter as a random effect for controls. Only results from models w/ litter as a random effect are shown here. Regions overlapping with previous significant analysis results are noted by timepoint. KW and BFL here refers to the univariate analysis (see section 3.1.1).

| Model w/ and w/o litter as a random effect AIC comparisons |  |  |  |  |  |  |  |  |  |
| --- | --- | --- | --- | --- | --- | --- | --- | --- | --- |
| Model | K | AICc | AICcWt | LL | Region | PND |  |  |  |
|  |  |  |  |  |  | 21 | 38 | 60 | 90 |
| model_3 | 6 | 253.13 | 0.49 | -120.24 | CA_L |  |  | KW |  |
| model_3 | 6 | 285.83 | 0.41 | -136.59 | STRv_L | KW<br>PC2 |  |  |  |
| model_3 | 6 | 301.47 | 0.57 | -144.41 | STRv_R |  |  |  |  |
| model_3 | 6 | 305.63 | 0.48 | -146.49 | PIR_R | PC2 |  | KW |  |
| model_3 | 6 | 337.97 | 0.39 | -162.66 | PIR_L | PC2 |  |  |  |
| model_3 | 6 | 365.28 | 0.58 | -176.31 | PERI_L |  |  | KW |  |
| model_3 | 6 | 383.91 | 0.49 | -185.63 | SSp_L |  | BFL | PC2 | KW<br>PC3<br>LV2 |
| model_4 | 7 | 303.45 | 0.65 | -144.28 | FRP_L | PC2 |  |  |  |
| model_4 | 7 | 321.01 | 0.67 | -153.06 | VERM_L |  |  |  |  |
| model_4 | 7 | 342.40 | 0.52 | -163.76 | AUD_L | PC2 |  |  |  |
| model_4 | 7 | 347.11 | 0.78 | -166.11 | COA_R |  |  |  | PC3<br>LV2 |
| model_4 | 7 | 363.44 | 0.60 | -174.28 | ECT_L | KW<br>PC2 |  |  |  |
| model_4 | 7 | 371.32 | 0.44 | -178.22 | SSp_R |  |  |  | KW<br>PC2<br>PC3<br>LV2 |
| model_4 | 7 | 373.39 | 0.40 | -179.25 | TEa_L | PC2 |  |  | PC3<br>LV2 |
| model_4 | 7 | 377.88 | 0.62 | -181.50 | SSs_L |  |  |  | PC3<br>LV2 |

AICc: Akaike Information Criterion corrected, k: number of estimated parameters in the model, AICcWt: Akaike weights, LL: Log-likelihood.

**Table 2 :** Examples of brain measures LMER estimates for models w/ and w/o litter as a random effect. Models inputs: z-scored structure volumes and age, sex [male, female; referencing male], mouse id and litter as factors.

| LMER estimates for models w/ and w/o litter as a random effect |  |  |  |  |  |  |  |
| --- | --- | --- | --- | --- | --- | --- | --- |
| Region | Model | Ranef<br>Litter | sex<br>[effect size<br>(p-val)] | age<br>[effect size<br>(p-val)] | sex*age<br>[effect size<br>(p-val)] | litter<br>[var, s.d.] | mouse id<br>[var, s.d.] |
| COA_R | model_4 | w/ | 0.184<br>(0.249) | 0.300<br>(0.002) | 0.304<br>(0.029) | 0.127,<br>0.356 | 0.000,<br>0.000 |
|  | model_2 | w/o | 0.099<br>(0.581) | 0.298<br>(0.003) | 0.310<br>(0.031) | NA | 0.106,<br>0.325 |
| PIR_L | model_3 | w/ | 0.021<br>(0.908) | 0.540<br>(0.000) | NA | 0.066,<br>0.257 | 0.079,<br>0.281 |
|  | model_1 | w/o | 0.025<br>(0.890) | 0.540<br>(0.000) | NA | NA | 0.137,<br>0.371 |
| CA_L | model_3 | w/ | 0.339<br>(0.032) | 0.745<br>(0.000) | NA | 0.065,<br>0.256 | 0.093,<br>0.304 |
|  | model_1 | w/o | 0.315<br>(0.057) | 0.745<br>(0.000) | NA | NA | 0.159,<br>0.399 |

Ranef = Random effect, var. = variance, s.d.= Standard Deviation., w/ and w/o = with and without, NA = Not applicable

#### 3.2.2 Power simulation

Power simulations based on LMER models, including the litter as a random effect (normative development: model 3; group differences in development: model 5, see section 2.5.1) in an attempt to get maximum characterization of the litter-effect when present, were performed. For power simulations relative to every brain structure modeling normative development, 10 regions reached 80% power for a 0.10 age effect size (the left and right visual area, Ammon’s horm of the hippocampus, and hindbrain, the left ventral striatum, striatum-like amygdalar nuclei, agranular insular/orbital areas, and the right ectorhinal cortex), 69 for a 0.15 effect size (except for the left anterior cingulate cortex, caudal region of the pallidum and the right piriform-amygdalar area), and all regions (72) for 0.20 and 0.25 effect sizes. From sample size estimates for every brain region, SS and MM measures were extracted. When comparing the MM and SS results, the best sample size (SS) was achieved with a greater number of litters, fewer mice per litter (< 6), and a smaller overall sample size (Table 4) for 48 and 34 brain regions for an age effect of 0.20 (Table 3) and 0.25 respectively. A total of 21 brain regions were shared and significant for both effect sizes (Supplementary Table S6). These regions benefited from having an optimized sample size where the number of litters is greater. Regions such as the primary somatosensory area and the ventral striatum needed a much smaller sample size to observe an age effect compared to the anterior olfactory bulb, the anterior cingulate cortex and the pallidum (Table 3, Fig. 10).

**Fig. 10 :**
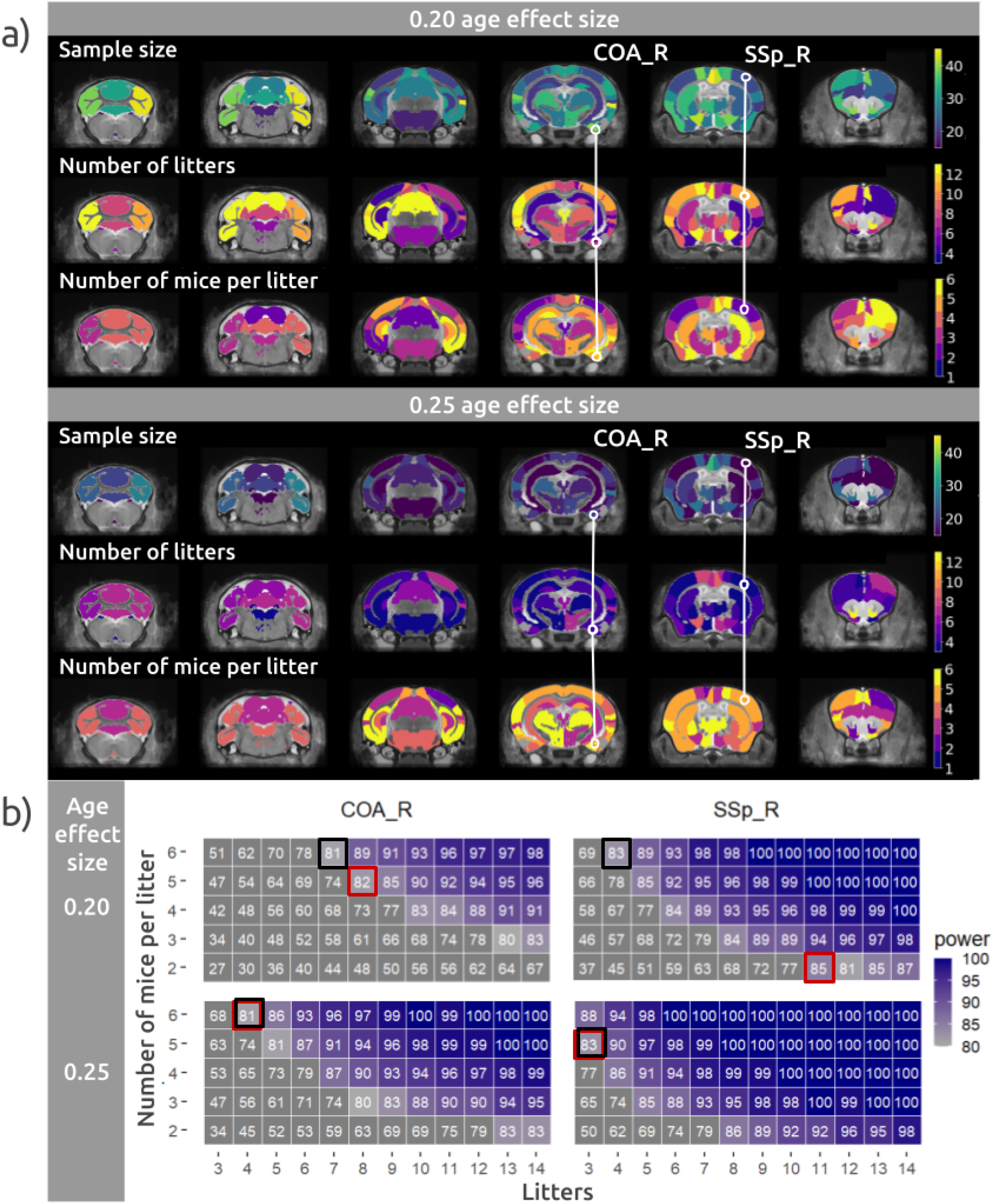
Power simulation results for normative development modeling. Results from an 0.20 and 0.25 age effect size are presented. a) Brain maps displaying sample size, number-of-litters and number of mice per litter (litter-size) for best SS trade-off results. b) Power estimate heatmaps for selected brain regions. Red square = SS trade-off, Black square = MM trade-off. COA_R = Right cortical-amygdalar region, SSp_R = Right Primary somatosensory area.

**Table 3 :** Regions with SS achieving a smaller sample size compared to MM for normative development modeling with a 0.20 age effect size. See **Supplementary Table S2** for regions acronyms. (Pink = Regions highlighted by the AIC analysis (Table 1)).

| Best sample size trade-offs from power simulation based on normative development |  |  |  |  |  |  |  |
| --- | --- | --- | --- | --- | --- | --- | --- |
| Effect | Region<br>(48/72) | Number<br>of<br>litters |  | Number<br>of<br>mice per litter |  | Sample size |  |
|  |  |  |  | SS(MM) | SS-MM |  |  |
| 0.20 | ACA_L | 13(10) | 3 | 4(6) | -2 | 52(60) | -8 |
| 0.20 | SSs_L | 10(6) | 4 | 3(6) | -3 | 30(36) | -6 |
| 0.20 | PTLp_L | 10(6) | 4 | 3(6) | -3 | 30(36) | -6 |
| 0.20 | ECT_L | 12(7) | 5 | 3(6) | -3 | 36(42) | -6 |
| 0.20 | PIR_L | 9(7) | 2 | 4(6) | -2 | 36(42) | -6 |
| 0.20 | PALv_L | 14(8) | 6 | 3(6) | -3 | 42(48) | -6 |
| 0.20 | AUD_L | 13(5) | 8 | 2(6) | -4 | 26(30) | -4 |
| 0.20 | AI-ORB_L | 5(4) | 1 | 4(6) | -2 | 20(24) | -4 |
| 0.20 | RSP_R | 8(6) | 2 | 4(6) | -2 | 32(36) | -4 |
| 0.20 | RSP_L | 8(6) | 2 | 4(6) | -2 | 32(36) | -4 |
| 0.20 | PERI_L | 13(5) | 8 | 2(6) | -4 | 26(30) | -4 |
| 0.20 | DG_R | 8(6) | 2 | 4(6) | -2 | 32(36) | -4 |
| 0.20 | RHP_L | 13(5) | 8 | 2(6) | -4 | 26(30) | -4 |
| 0.20 | PIR_R | 8(6) | 2 | 4(6) | -2 | 32(36) | -4 |
| 0.20 | COA_L | 8(6) | 2 | 4(6) | -2 | 32(36) | -4 |
| 0.20 | sAMY_R | 13(5) | 8 | 2(6) | -4 | 26(30) | -4 |
| 0.20 | VERM_R | 8(6) | 2 | 4(6) | -2 | 32(36) | -4 |
| 0.20 | VERM_L | 8(6) | 2 | 4(6) | -2 | 32(36) | -4 |
| 0.20 | HEM_R | 11(8) | 3 | 4(6) | -2 | 44(48) | -4 |
| 0.20 | MOs_L | 9(5) | 4 | 3(6) | -3 | 27(30) | -3 |
| 0.20 | FRP_L | 11(6) | 5 | 3(6) | -3 | 33(36) | -3 |
| 0.20 | AUD_R | 11(6) | 5 | 3(6) | -3 | 33(36) | -3 |
| 0.20 | VIS_R | 7(4) | 3 | 3(6) | -3 | 21(24) | -3 |
| 0.20 | ILA-PL_R | 13(7) | 6 | 3(6) | -3 | 39(42) | -3 |
| 0.20 | TEa_L | 13(7) | 6 | 3(6) | -3 | 39(42) | -3 |
| 0.20 | PERI_R | 9(8) | 1 | 5(6) | -1 | 45(48) | -3 |
| 0.20 | AON_R | 13(7) | 6 | 3(6) | -3 | 39(42) | -3 |
| 0.20 | PALd_L | 9(5) | 4 | 3(6) | -3 | 27(30) | -3 |
| 0.20 | PALv_R | 13(7) | 6 | 3(6) | -3 | 39(42) | -3 |
| 0.20 | TH_R | 9(5) | 4 | 3(6) | -3 | 27(30) | -3 |
| 0.20 | HY_L | 9(5) | 4 | 3(6) | -3 | 27(30) | -3 |
| 0.20 | HEM_L | 13(7) | 6 | 3(6) | -3 | 39(42) | -3 |
| 0.20 | SSs_R | 7(5) | 2 | 4(6) | -2 | 28(30) | -2 |
| 0.20 | SSp_R | 11(4) | 7 | 2(6) | -4 | 22(24) | -2 |
| 0.20 | SSp_L | 11(4) | 7 | 2(6) | -4 | 22(24) | -2 |
| 0.20 | TEa_R | 7(5) | 2 | 4(6) | -2 | 28(30) | -2 |
| 0.20 | AON_L | 8(7) | 1 | 5(6) | -1 | 40(42) | -2 |
| 0.20 | COA_R | 8(7) | 1 | 5(6) | -1 | 40(42) | -2 |
| 0.20 | STRv_R | 7(5) | 2 | 4(6) | -2 | 28(30) | -2 |
| 0.20 | PALc_R | 8(7) | 1 | 5(6) | -1 | 40(42) | -2 |
| 0.20 | MB_L | 14(5) | 9 | 2(6) | -4 | 28(30) | -2 |
| 0.20 | HB_L | 6(4) | 2 | 3(5) | -2 | 18(20) | -2 |
| 0.20 | MOp_L | 13(8) | 5 | 3(5) | -2 | 39(40) | -1 |
| 0.20 | MOs_R | 7(6) | 1 | 5(6) | -1 | 35(36) | -1 |
| 0.20 | STRd_L | 7(6) | 1 | 5(6) | -1 | 35(36) | -1 |
| 0.20 | PALd_R | 8(5) | 3 | 3(5) | -2 | 24(25) | -1 |
| 0.20 | TH_L | 7(6) | 1 | 5(6) | -1 | 35(36) | -1 |
| 0.20 | HY_R | 7(6) | 1 | 5(6) | -1 | 35(36) | -1 |

Pink highlight = brain regions overlap between significant LMER and power simulation results.

**Table 4 :** Total sample size reference table for each number of mice-per-litter (litter-size) to number-of-litters combination. The 24 sample size is highlighted only as a reference. No assumptions are made here regarding to mice sex.

**Total sample size for litter and mice per litter combination**
|  |  |  |  |  |  |  |  |  |  |  |
| --- | --- | --- | --- | --- | --- | --- | --- | --- | --- | --- |
| 6 | 18 | 24 | 30 | 36 | 42 | 48 | 54 | 60 | 66 | 72 |
| 5 | 15 | 20 | 25 | 30 | 35 | 40 | 45 | 50 | 55 | 60 |
| 4 | 12 | 16 | 20 | 24 | 28 | 32 | 36 | 40 | 44 | 48 |
| 3 | 9 | 12 | 15 | 18 | 21 | 24 | 27 | 30 | 33 | 36 |
| 2 | 6 | 8 | 10 | 12 | 14 | 16 | 18 | 20 | 22 | 24 |
| Mice↑ | 3 | 4 | 5 | 6 | 7 | 8 | 9 | 10 | 11 | 12 |
| Litters |  |  |  |  |  |  |  |  |  |  |

For power simulations modeling simulated treatment group differences, 50 regions for a 0.35 treatment group by age effect size (data not shown) reached 80% power, and all regions (72) for 0.40 and 0.45 effect sizes. The regional relationship for sample sizes were maintained relative to the previous analysis. For example, the hippocampus ammon’s horn needed a smaller sample size and the anterior cingulate cortex a bigger sample size (Fig. 11). When comparing SS and MM results, 32 regions reached 80% with smaller sample sizes for SS trade-offs for a 0.40 treatment group by age effect size, 33 regions for a 0.45 effect size, and 11 regions overlapped between both effect sizes (Supplementary Table S6). From the regions selected in **Table 3** and **5**, regions differences and overlaps were seen in the SS and MM comparisons, between normative development and treatment group power simulation (Supplementary Table S6).

**Fig. 11 :**
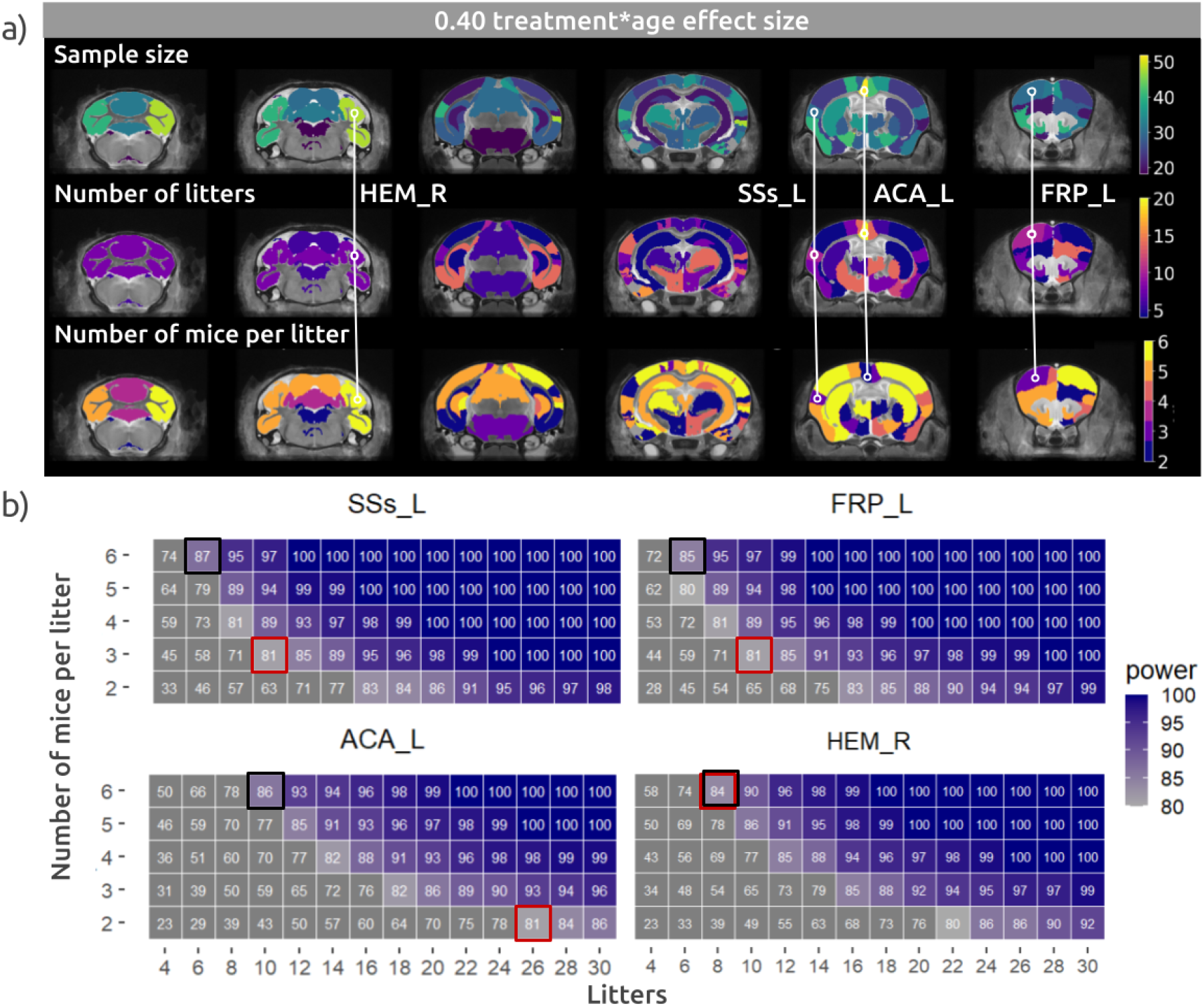
Power simulation results for treatment group differences modeling. Results from a 0.40 treatment group by age effect size. a) Brain maps displaying sample size, number-of-litters and number of mice per litter (litter-size) for best SS trade-off results. b) Power estimates heatmaps for selected brain regions. Red square = SS trade-off, Black square = MM trade-off. SSs_L = Left Supplemental somatosensory area, FRP_L = Left Frontal pole, ACA_L = Left Anterior cingulate cortex, HEM_R = Right cerebellar Hemispheric regions,.

**Table 5 :**
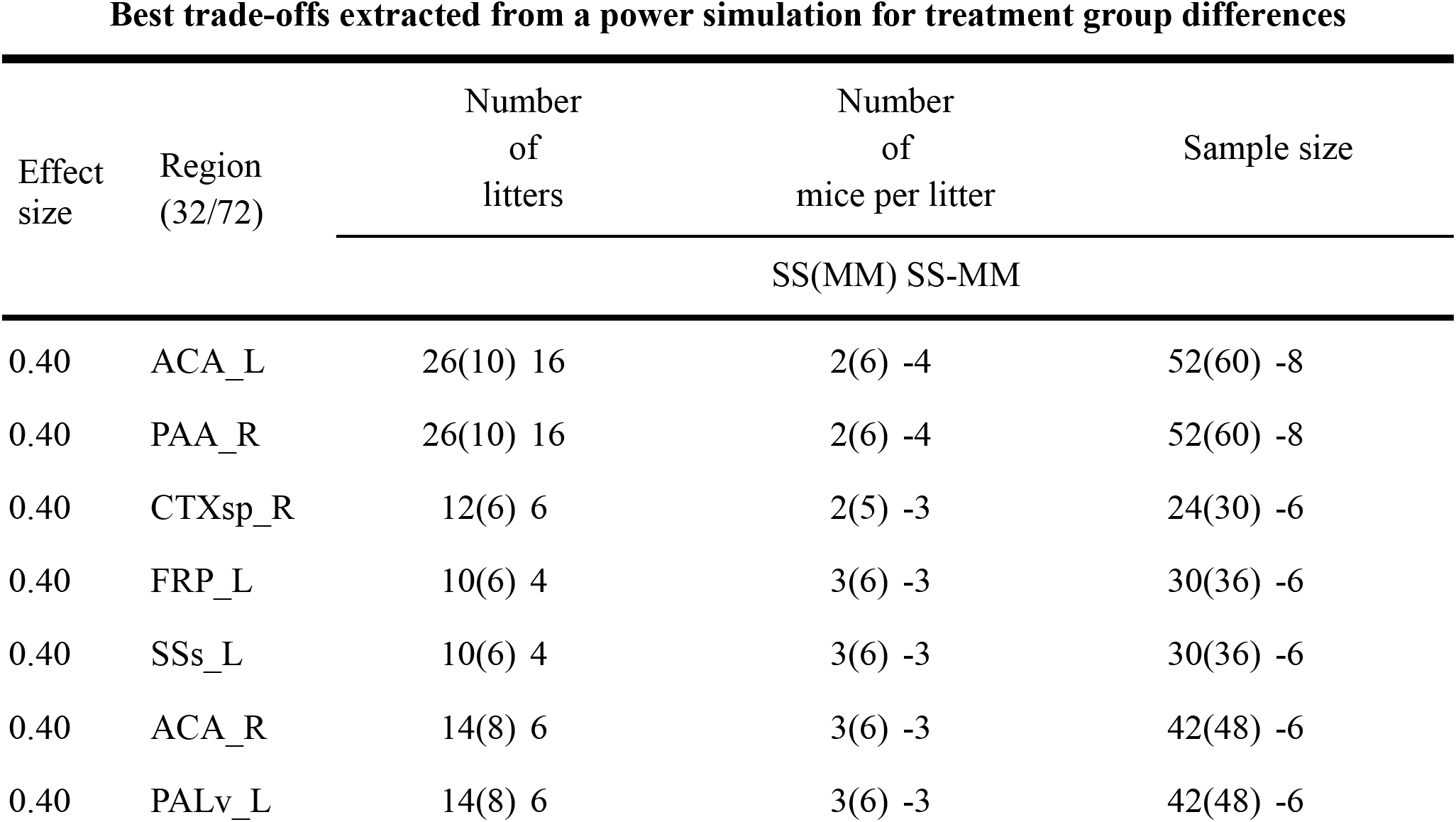
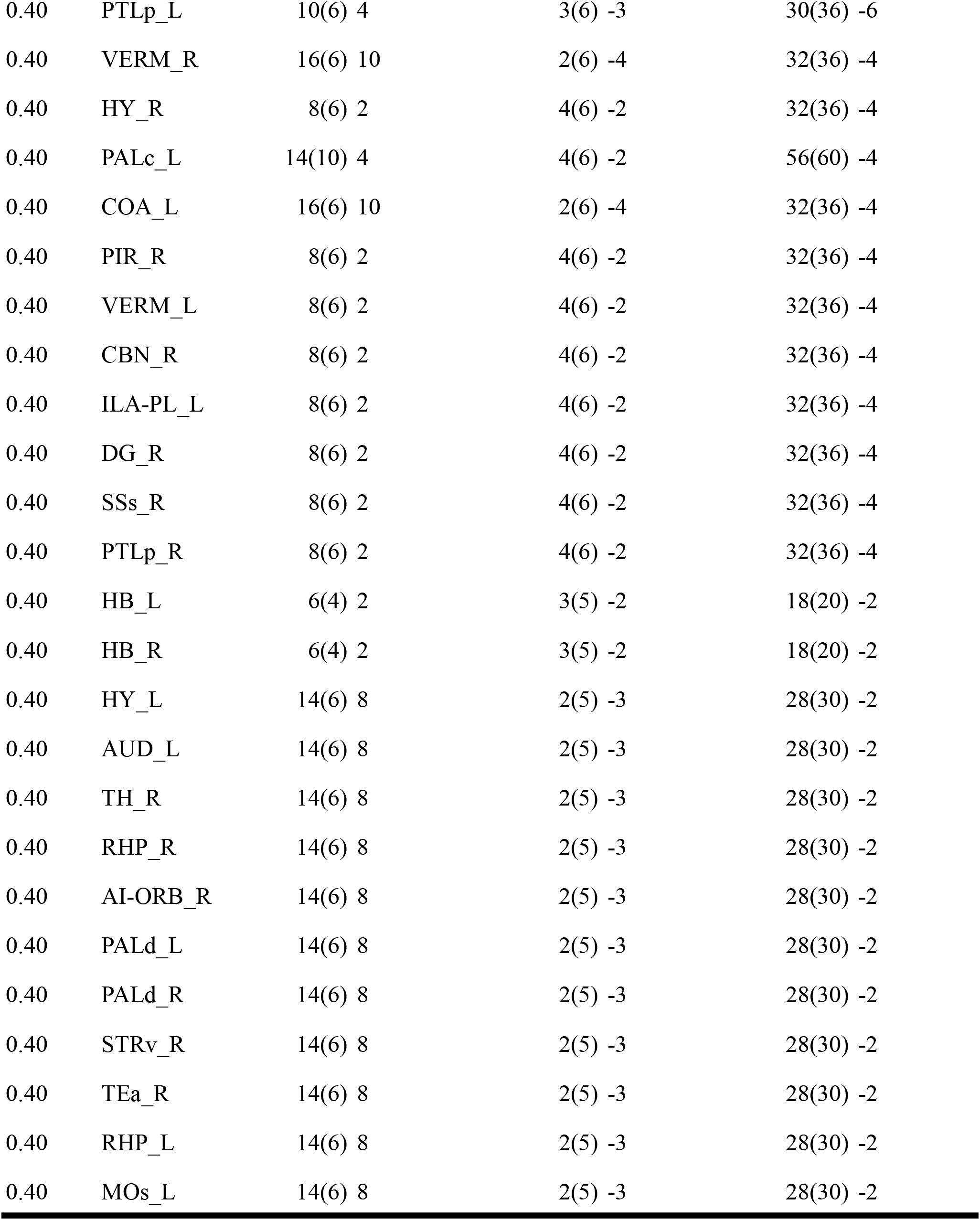
Regions with SS achieving a smaller sample size compared to MM for treatment group differences modeling with a 0.40 treatment*age effect size. See **Supplementary Table S2** for regions acronyms.

| Best trade-offs extracted from a power simulation for treatment group differences |  |  |  |  |
| --- | --- | --- | --- | --- |
| Effect size | Region (32/72) | Number of litters | Number of mice per litter | Sample size |
|  |  | SS(MM) SS-MM |  |  |
| 0.40 | ACA_L | 26(10) 16 | 2(6) -4 | 52(60) -8 |
| 0.40 | PAA_R | 26(10) 16 | 2(6) -4 | 52(60) -8 |
| 0.40 | CTXsp_R | 12(6) 6 | 2(5) -3 | 24(30) -6 |
| 0.40 | FRP_L | 10(6) 4 | 3(6) -3 | 30(36) -6 |
| 0.40 | SSs_L | 10(6) 4 | 3(6) -3 | 30(36) -6 |
| 0.40 | ACA_R | 14(8) 6 | 3(6) -3 | 42(48) -6 |
| 0.40 | PALv_L | 14(8) 6 | 3(6) -3 | 42(48) -6 |
| 0.40 | PTLp_L | 10(6) 4 | 3(6) -3 | 30(36) -6 |
| 0.40 | VERM_R | 16(6) 10 | 2(6) -4 | 32(36) -4 |
| 0.40 | HY_R | 8(6) 2 | 4(6) -2 | 32(36) -4 |
| 0.40 | PALc_L | 14(10) 4 | 4(6) -2 | 56(60) -4 |
| 0.40 | COA_L | 16(6) 10 | 2(6) -4 | 32(36) -4 |
| 0.40 | PIR_R | 8(6) 2 | 4(6) -2 | 32(36) -4 |
| 0.40 | VERM_L | 8(6) 2 | 4(6) -2 | 32(36) -4 |
| 0.40 | CBN_R | 8(6) 2 | 4(6) -2 | 32(36) -4 |
| 0.40 | ILA-PL_L | 8(6) 2 | 4(6) -2 | 32(36) -4 |
| 0.40 | DG_R | 8(6) 2 | 4(6) -2 | 32(36) -4 |
| 0.40 | SSs_R | 8(6) 2 | 4(6) -2 | 32(36) -4 |
| 0.40 | PTLp_R | 8(6) 2 | 4(6) -2 | 32(36) -4 |
| 0.40 | HB_L | 6(4) 2 | 3(5) -2 | 18(20) -2 |
| 0.40 | HB_R | 6(4) 2 | 3(5) -2 | 18(20) -2 |
| 0.40 | HY_L | 14(6) 8 | 2(5) -3 | 28(30) -2 |
| 0.40 | AUD_L | 14(6) 8 | 2(5) -3 | 28(30) -2 |
| 0.40 | TH_R | 14(6) 8 | 2(5) -3 | 28(30) -2 |
| 0.40 | RHP_R | 14(6) 8 | 2(5) -3 | 28(30) -2 |
| 0.40 | AI-ORB_R | 14(6) 8 | 2(5) -3 | 28(30) -2 |
| 0.40 | PALd_L | 14(6) 8 | 2(5) -3 | 28(30) -2 |
| 0.40 | PALd_R | 14(6) 8 | 2(5) -3 | 28(30) -2 |
| 0.40 | STRv_R | 14(6) 8 | 2(5) -3 | 28(30) -2 |
| 0.40 | TEa_R | 14(6) 8 | 2(5) -3 | 28(30) -2 |
| 0.40 | RHP_L | 14(6) 8 | 2(5) -3 | 28(30) -2 |
| 0.40 | MOs_L | 14(6) 8 | 2(5) -3 | 28(30) -2 |

A tendency was observed across a considerable number of regions, about half of the brain regions, such as to reach a 80% power threshold with the smallest sample size, a greater number of litters but of smaller size were needed. This seems to further support the idea of a litter-effect, but specific to some brain regions and type/size of effects (age effect size, treatment group by age effect size) when modeling brain volumetric measures development.

## 4. Discussion

In this manuscript we analyzed the impact of litter effects in mouse development by examining litter differences in brain structure volumetry (MAGeT brain structures’ extracted volumes) and behavioral measures (MBT, PPI, and PPI) across the lifespan. This was evaluated at the univariate level for each measure independently and at the multivariate level identifying shared patterns across brain structures, and their shared covariance with behaviors. Pertaining to the litter, we observed greater variance in distinct measures in the adolescent and adult period and more integrative patterns of brain and behaviors in adulthood, but not in adolescence. Finally, we provided a comprehensive set of power analysis, modeling brain change and group differences to better inform future design decisions related to sample size and litter variation.

### 4.1 Normative development

#### 4.1.1 Litter-specific development

We first wanted to evaluate if we could observe brain and behavioral differences within or between litters in normative mouse development, and if so, were there specific brain patterns associated and would they covary with behavioral patterns. Initially, a greater variance of measures between than within litter was predicted. In the childhood period (PND 21), we observed litters difference in their associations to patterns (PC2) of bigger regions related to olfaction and emotional regulation (cortical-, piriform-amygdalar, and piriform areas), and smaller regions of higher order of integration (temporal association, central striatum, frontal pole, auditory, ectorhinal cortices). Meaning mice from the same litter associated to these pattern in a similar fashion compared to mice from a different litter. In other words, we did observe specific measures varying and grouping by litter. More specifically, some regions highlighted by our results, for being modulated by a litter-effect, also associate with developmental or group associate specificities such as hierarchical cortical maturation, social behaviors and sex dimorphism. For example, the temporal association and ectorhinal areas are related to the development of higher visual processing (Nishio et al., 2018), the amygdalar regions are sexually dimorphic and develop early (Brenhouse & Andersen, 2011; Premachandran et al., 2020) and their association to frontal areas linked to social behaviors (Bicks et al., 2015). These regions associated with this specific principal component seem to show a hierarchical patterning of cortical maturation (Chomiak & Hu, 2017), where a relationship of smaller higher integration regions and bigger primary sensory or subcortical regions was observed. These results might highlight some form of litter-dependant neurodevelopment trajectories for these specific regions.

Modeling volumetric developmental trajectories (LMERs) across brain regions was improved by the addition of a random effect of litter for some regions. The improved models associated to specific regions were also consistent with regions highlighted by volumetric patterns (PCs), either in the early developmental period and in adulthood (Table 1). It could possibly highlight an early emergence of the litter-effect in childhood that becomes difficult to detect during adolescence due to asynchronous maturational changes between mice (e.g. sex neurodevelopmental differences, experience differences), and detectable again later in adulthood.

Individualization, the conceptual process of becoming a unique individual, could be modulated by enduring litter-effect as a result of postnatal environmental impact (Freund et al., 2013; Lathe, 2004). However, from our results, covariance of brain and behavioral measures have only been seen in adulthood. We could hypothesize that a litter-effect might arise in early development and be attributable to individual factors relative to a litter (e.g. univariate results) and grow to be greater and/or possibly more complex throughout development exhibited by broad changes in brain patterns and their association to behaviors (multivariate results).

There have been considerable arguments supporting greater brain sensitivity to enriched and challenging environments and biological variability during adolescence (Marco et al., 2011). The heterogeneity in litter medians and variances was observed for the left thalamus in the early adolescence period (PND 38) in co-occurrence with variations of behavioral measures associated with sensorimotor and locomotor activity (OFT). In later adolescence (PND 60), no behavioral differences were observed, but some regions (dorsal striatum, ammon’s horn, piriform area, hemispheric cerebellar regions) (Fig. 5 and Fig. 6, Supplementary Table S4 and Table S5) showed litter-specific differences at a subthreshold level. These regions have been reported to be highly sensitive to the environment (Marco et al., 2011; Szulc et al., 2015).

We believe that different factors, such as the pup sex (Premachandran et al., 2020; Qiu et al., 2018), which may or may not pertain to the litter environment seem to create greater variance during adolescence and early adutlhood. In adolescence no relevant integrative measures (PCA, PLSC) such as brain patterns or their covariance with behavioral measures were observed, but significant brain patterns were in early adulthood. This may be due to an overlap or a heterosynchronous sex specific brain growth trajectories (Qiu et al., 2018), a litter-effect, and other environmental effects.

Anxiety-like, risk-taking and sensorimotor gating responses have previously been observed to go through dynamic changes during the adolescence period (Fairless et al., 2012; Laviola et al., 2003; Marco et al., 2011). The most relevant brain and behavioral patterns were seen right after these changes, in the adulthood period (PND 90) demonstrating covariation patterns between sensory integration and motor commands regions, and exploratory behaviors (OFT: center measures), varying in opposite directions to memory, cerebellar and emotion associated regions, and anxiety-like behaviors (OFT: corners measures).

#### 4.1.2 Social behaviors development

Many of the brain regions highlighted wihtin this project, based on all analyses results, are highly susceptible to the environment (Marco et al., 2011; Szulc et al., 2015) and could represent a juxtaposition of litter- and sex-effects linked to memory, social behaviors (Cooke et al., 2007) and social recognition (Ko, 2017; Raam & Hong, 2021; Tan et al., 2019). More precisely, medial temporal areas (hippocampus, amygdala, perirhinal area, etc), frontal, and cerebellar regions have important roles in social recognition via olfaction (Camats Perna & Engelmann, 2017), memory (Srinivasan & Stevens, 2018), and social attachment patterns (Coria-Avila et al., 2014). These regions are part of functional regional circuits supporting these behaviors. Furthermore, the subcortical regions follow a sexually dimorphic trajectory (Fairless et al., 2012; Premachandran et al., 2020). Also supporting this, the adolescence is known to be a sensitive period for the thalamo-PFC circuit maturaturation (O’Reilly et al., 2021) and HPA-axis development (Schmidt et al., 2003), associated with social behaviors and fear respectively. Within-litter environment is also favorable to social hierarchy, especially when the space is shared between males and females. It is a common organization seen in groups of animals, such as mice, and it establishes roles and social status (Nelson et al., 2019; Zhou et al., 2017). Furthermore, social behaviors are known to be significantly impaired in neurodevelopmental disorders. In autism spectrum disorder for example, abnormal connection between the prefrontal cortex and amygdala has been shown to be related to social impairment (Huang et al., 2016). From our current results, a litter-effect could impact the observed phenotypes also studied in animal model studies of neurodevelopmental disorders.

### 4.2 Power simulation of normative and treatment-like mice development

The need for well-powered studies has been a long-lasting discussion in research and should be standard practice prior to designing studies. Statistical power calculation tools are broadly available, estimating trade-offs between the desired effect size, sample size, variance and significance level. Our results demonstrated a need for different sample sizes, made of different litter to mice-per-litter ratios, for each individual region when observing specific effects. Regions previously associated, within the scope of our analysis, to a possible litter-effect were part of the regions that benefited from a larger number of litters instead of a greater number of mice per litter. Taken together, when modelling normative development, 15 regions benefited from a model including litter as a random effect. A total of 48 out of 72 regions, including 12 of the 15 regions from the LMER analysis, did reach 80% power with a smaller sample size when having more litters without maximizing the number of mice per litter. When modeling treatment-like group differences, 38 regions did reach the same consensus as the normative development model. These results could possibly underline a litter-effect in some regions, in this case representing close to half of the brain, where a higher ratio of litters is needed and is imperative to achieve high power. It also emphasizes the heterogeneous developmental trajectories of brain structures (Narvacan et al., 2017; Raznahan et al., 2013), by the range of effect sizes and how probable they could be detected related to specific sample size characteristics.

### 4.3 Limitations and future perspectives

We do recognize some limitations within this study pertaining to different factors such as methodology choices and sample availability. First, standardized procedures and protocols were used in an aim to maintain consistency between assessments, but are a great limiting factor to the generalization of results. Second, our analysis was done on a small sample size extracted from a previous study sample and acquired for a different purpose then what we intended to observe. A dataset with a greater focus on this litter-effect could benefit from more targeted behavioral assessments. Third, we modeled growth trajectory with simple LMER models but the use of a more complex spline fit (i.e. Cubic, Quadratic, Linear) for individual brain region changes should be explored in the future (Shaw et al., 2008). Fourth, we evaluated gross anatomy associated with the volumes of fewer brain areas since this project was exploratory. Evaluating the litter-effect in more details with multi-modal data and analysis at the voxel level would be of great interest, in light of the present findings. Furthermore, to make the best use of the longitudinal data format, future multivariate analysis with LMER effect sizes from these different measures as inputs could be explored. Fifth, our power simulation only focused on the main effects we wanted to observe such as the age and group (treatment) effect and a ceiling of 6 mice per litter was selected. The exploration of a greater array of effects and varying ceiling numbers would be of interest. Effects such as sex for example, would be a crucial future inverstigation because of its potential interaction with the litter-effect.

The repeated discussions about the reproducibility crisis and translational issues related to research using preclinical/animal models has resulted in an interest in standardization and guidelines. Future neurodevelopmental designs such as cross-fostering (McCarty, 2017), a model where a dam fosters pups carried by another dam, or systematic heterogeneity, a research model varying multiple factors in a systematic fashion, highlighted by different groups in the past couple of years (Brubaker et al., 2019; Goot et al., 2021; Richter, 2017; Richter et al., 2011; Weber-Stadlbauer & Meyer, 2019) might be a great path of exploration for improving litter standardization. With the innovative impact of computer science within neuroscience, this new symbiosis can now lead to the use of more complex statistical tools. With the aggregation of data from numerous locations, tools such as ComBat and CovBat have been developed to harmonize data variation by correcting for bacth effects, for example from different acquisition sites (Chen et al., 2022). At a smaller scale, harmonization of data between litters could be explored using adapted versions of these tools.

### 4.4 Guidelines

In light of these results and previous guidelines regarding neurodevelopmental studies, here are some conclusions and recommendations:

1. Longitudinal assays of brain anatomy and behaviors in mice may be variable and impacted by the litter. This manuscript provides a guide for the characterization of this variability. While it remains to be seen how well these results may generalize across research groups, spectrum of MRI acquisitions or behavioral tests, the methods proposed are readily available and commonly implemented in the neuroimaging literature.
2. In longitudinal volumetrics studies, the investigator should consider the specific regions where they expect to observe variation in the rate of anatomical change. Power estimates that we have demonstrated in this manuscript suggest that there are specific design choices that should be made with respect to the trade-off between number-of-litters, litter-size and overall sample size that impact the study design.
3. Investigators should consider performing a similar characterization for the MRI and behavioral measures that they preferentially use, as these methods vary greatly between laboratories. Better characterization may provide further methods for defining sample size estimates while giving appropriate consideration to the litter effect.
4. Sex effects may be modulated by the litter via sex-ratios within litters. Since this variable is litter-specific, a random effect of litter should be included in mixed effects models as it is crucial for accounting for this dual source of variability.
5. Our results strongly suggest that litter membership impacts brain regions that have a role in social behaviors processing and should be further accounted for when examining these classes of behaviors.

## 5. Conclusion

In conclusion, greater variance of measures was seen in the adolescence period and a litter-effect was observed in brain, behavior and their relationships in normative developing mice throughout the lifespan. As suggested in previous studies, greater similarities of neurobiologic and behavioral measures between mice from the same litters were expected (Golub & Sobin, 2020). According to our results, only specific brain regions and patterns are impacted by this litter-effect. From our brain and behavior analyses, patterns of cerebral regions related to the neural circuitry of social behavior and recognition could be modulated by the litter, and so could their covariance with exploratory behaviors. As observed in our power analysis, to maintain a small sample size and achieve high power, a greater number of litters is required when assessing differences (rate of neurodevelopment within mice or treatment-like group differences) across a large number of brain regions sparsly distributed in space. This litter-effect also seems to be present throughout development and living in the shadow of significant sex-effects (Vorhees & Williams, 2021). These results will have repercussions on research methods decisions in studies modeling neurodevelopmental disorders (e.g. autism spectrum disorder, schizophrenia) and further enhance reproducibility. As supported by recent guidelines (Golub & Sobin, 2020), (Jiménez & Zylka, 2021), and (Vorhees & Williams, 2021), there is a pressing need to encourage the control of the litter in preclinical studies, and especially the one focusing on development. The control of this litter-effect could, in part, correct for different factors such as the impact of maternal care, litter size and in utero and postnatal environment intrinsically linked to the litter. These results confirm an observable litter-effect and lead the way to the expansion of this knowledge in the future.

## Supporting information

Supplementary methods and results

## List of Abbreviations

AIC: akaike information criterion
BFL: brown-forsynthe-levene test
dB: decibels
EE: environmental enrichment
FDR: false discovery rate
GD: gestational day
HPA-axis: hypothalamus-pituitary-adrenal axis
KW: kruskal-wallis test
PC: principal components
PCA: principal component analysis
PLSC: partial least squares correlation
PND: postnatal day
LMER: linear mixed effects model
LV: latent variable
MAGeT: multiple automatically generated templates
MBT: marble burying task
MIA: maternal immune activation
MM: maximum number of mice per litter
MnCl_2_: manganese chloride
MRI: magnetic resonance imaging
OFT: open field test
PPI: prepulse inhibition test
SAL: saline
SS: smallest sample size

