## Supplementary methods and results for "Examining Litter Specific Variability in Mice and its Impact on Neurodevelopmental Studies"

### Supplementary methods

**Method S1:** Behavioral testing description

**Marble burying task** : Mice were placed in a home cage filled with bedding (~7cm of woodchip) where 15 marbles were placed at equal distances. After 30 minutes (video recorded), marbles were classified either as 100% buried (not visible), 75% buried (slightly visible) or unburied (<75% buried).

**Prepulse inhibition test**: Mice were placed in a sound-attenuating plexiglass chamber (video recorded), where a speaker was mounted to provide background noise (70 dB) and acoustic stimuli, and a motion detector present to measure the mouse startle response. The startle response was measured (startle tone 120 dB), without prepulse, for the first 8 trials and final 7 trials. For the remaining 35 trials, the startle response either from the startle tone alone, or the prepulse response from the startle tone preceded by a random prepulse stimuli of 3, 6, 9, 12 or 15dB above background noise, for a total of 5 each, were measured. Percent PPI from each prepulse intensity (averaged over trials) was calculated : [(startle response - prepulse response) / startle response] x 100.

**Open field test** : Mice were placed in an arena of 45 x 45 cm (video recorded), where different zones (center, edges, corners) were conceptually delimited. During the 15 minutes where the mouse explored the arena, the distance, frequency and time spent in these different zones were measured.

N.B. All videos were later analyzed using Ethovision XT12 (Noldus Information Tech Inc., Leesburg, VA) (Guma, do Couto Bordignon, et al., 2021).

**Table S1**: Detailed distribution of male and female (m:f) control pups assessed per litter and by timepoints (Guma et al., 2022).

**Litter and sex distribution for selected mice**

| | Litters | | | | | | |
|---|---|---|---|---|---|---|---|
| | 1 | 2 | 3 | 4 | 5 | 6 | 7 |
| PND21 | 4:2 | 3:3 | 1:4 | 0:3 | 2:1 | 0:3 | 4:0 |
| PND38 | 4:2 | 3:3 | 2:2 | 1:4 | 2:2 | 0:4 | 4:0 |
| PND60 | 4:2 | 4:3 | 2:4 | 1:4 | 2:2 | 0:4 | 4:0 |
| PND90 | 4:2 | 4:3 | 2:4 | 1:4 | 2:2 | 0:4 | 4:0 |
| Total | 4:2 | 4:3 | 2:4 | 1:4 | 2:2 | 0:4 | 4:0 |

**Table S2**: MAGeT brain 36 structures labels name.

**Brain structure labels**

| | | | |
|---|---|---|---|
| MOp | Primary motor area | CTXsp | Cortical subplate |
| MOs | Secondary motor area | AON | Anterior olfactory bulb |
| FRP | Frontal pole, cerebral cortex | PIR | Piriform area |
| SSs | Supplemental somatosensory area | COA | Cortical amygdalar area |
| SSp | Primary somatosensory area | PAA | Piriform-amygdalar area |
| AUD | Auditory areas | STRd | Striatum dorsal |
| VIS | Visual areas | STRv | Striatum ventral |
| ACA | Anterior cingulate cortex | sAMY | Striatum-like amygdalar nuclei |
| ILA-PL | Infralimbic/Prelimbic areas | PALd | Pallidum, dorsal region |
| AI-ORB | Agranular insular/orbital areas | PALv | Pallidum, ventral region |
| RSP | Retrosplenial area | PALc | Pallidum, caudal region |
| PTLp | Posterior parietal association areas | TH | Thalamus |
| TEa | Temporal association areas | HY | Hypothalamus |
| PERI | Perirhinal area | MB | Midbrain |
| ECT | Ectorhinal area | HB | Hindbrain |
| CA | Ammon's horn (Hipp) | VERM | Vermal regions |
| DG | Dentate gyrus (hipp) | HEM | Cerebellar Hemispheric regions |
| RHP | Retrohippocampal region | CBN | Cerebellar nuclei |

**Supplementary results**

**Table S3**: Shapiro-Wilk test assessing normality of measures distributions. Results showed significant non-normal distributions for some brain regions. Less regions were significant in earlier timepoints and more regions in adulthood.

| **Shapiro-Wilk test** | | | | |
|---|---|---|---|---|
| **Variables** | **PND21** | **PND38** | **PND60** | **PND90** |
| | **W, p-value** | | | |
| ACA_R | 0.920, 0.028 | | | |
| CTXsp_L | 0.921, 0.289 | | | |
| STRv_L | 0.925, 0.037 | | | |
| AON_R | 0.081, 0.003 | | | |
| HB_L | 0.927, 0.042 | | 0.925, 0.017 | |
| PIR_R | | 0.895, 0.004 | | |
| PIR_L | | 0.916, 0.015 | | |
| PAA_L | | 0.896, 0.004 | | |
| MOs_R | | 0.924, 0.024 | | |
| PTLp_R | | | 0.909, 0.006 | |
| VERM_R | | | 0.893, 0.002 | |
| HEM_R | | | 0.882, 0.001 | 0.888, 0.002 |
| SSp_R | | | 0.932, 0.030 | |
| AI-ORB_L | | | 0.924, 0.017 | |
| TH_L | | | 0.932, 0.029 | |
| MOs_L | | | | 0.918, 0.011 |
| FRP_R | | | | 0.909, 0.006 |
| VIS_L | | | | 0.935, 0.035 |
| ECT_L | | | | 0.926, 0.019 |
| COA_R | | | | 0.935, 0.035 |

| | | | | |
|---|---|---|---|---|
| STRd_L | | | | 0.850, 0.000 |
| sAMY_L | | | | 0.934, 0.033 |
| TH_R | | | | 0.819, 0.000 |
| MB_R | | | | 0.929, 0.024 |
| HEM_L | | | | 0.888, 0.002 |

**Table S4** : Brown-Forsythe-Levene test results for brain and behavioral measures. *From the distributions observations, it is likely that these results are driven by one outlier litter that exhibits an extreme variance.

**Brown-Forsythe-Levene test**

| **Variables** | **PND21** | **PND38** | **PND60** | **PND90** |
|---|---|---|---|---|
| | **F, p-value, q-value** | | | |
| SSp_L | | 4.563, 0.003, NA | | |
| PAA_R | | 3.797, 0.008, NA | | |
| TH_L | | 6.049, 0.000, 0.034 | | |
| Time_center | | 2.751, 0.032, NA | | |
| No_PP_AVG | | 4.103, 0.004, NA | | |
| AUD_R | | | 3.594, 0.009, NA | |
| AI-ORB_L | | | 5.924, 0.000, 0.034 | |
| PP_3dB_AVG | | | | 55.029, 0.000, 0.000* |
| PP_6dB_AVG | | | | 40.435, 0.000, 0.000* |
| PP_9dB_AVG | | | | 25.872, 0.000, 0.000* |
| PP_12dB_AVG | | | | 114.645,0.000,0.000* |
| PP_15dB_AVG | | | | 23.620, 0.000, 0.000* |

**Table S5**: Kruskal-Wallis test results for brain and behavioral measures at every timepoint.

| Variables | Kruskal-Wallis test<br>PND21 | PND38 | PND60 | PND90 |
|---|---|---|---|---|
| | H, p-value, q-value | | | |
| ECT_L | 18.179, 0.006, NA | | | |
| STRv_L | 17.560, 0.007, NA | | | |
| Distance corners | | 24.865, 0.000, 0.008 | | 15.843, 0.002, 0.014 |
| Frequency corners | | 24.145, 0.001, 0.008 | | 24.571, 0.000, 0.007 |
| Time corners | | 20.192, 0.003, 0.015 | | 21.411, 0.002, 0.014 |
| Distance edges | | 13.475, 0.036, NA | | 15.843, 0.015, 0.047 |
| Frequency edges | | 17.265, 0.002, 0.015 | | 25.110, 0.000, 0.007 |
| Time edges | | 21.058, 0.002, 0.041 | | 19.169, 0.004, 0.019 |
| Distance center | | 15.155, 0.019, NA | | 14.797, 0.022, NA |
| Frequency center | | 20.155, 0.003, 0.015 | | 19.767, 0.003, 0.017 |
| Distance total | | 23.260, 0.001, 0.008 | | 20.756, 0.002, 0.014 |
| PP_9dB_AVG | | 14.466, 0.025, NA | | 12.648, 0.049, NA |
| STRd_L | | | 20.501, 0.002, 0.041 | |
| STRd_R | | | 19.316, 0.004, 0.042 | |
| CA_L | | | 19.115, 0.004, 0.042 | |
| CA_R | | | 22.407, 0.001, 0.035 | |
| PIR_R | | | 25.750, 0.000, 0.018 | |
| HEM_L | | | 18.739, 0.005, 0.042 | |
| PERI_L | | | 19.031, 0.004, 0.042 | |
| SSp_L | | | | 20.310, 0.002, NA |
| SSp_R | | | | 19.698, 0.003, NA |
| CTXsp_L | | | | 17.462, 0.008, NA |
| PP_6dB_AVG | | | | 17.917, 0.006, 0.027 |

| | |
|---|---|
| PP_6dB_MAX | 5.527, 0.017, 0.047 |
| PP_12dB_AVG | 13.376, 0.037, NA |
| PP_15dB_AVG | 17.029, 0.009, 0.035 |
| MAX_startle_end | 12.677, 0.049, NA |
| AVG_startle_start | 14.211, 0.027, NA |

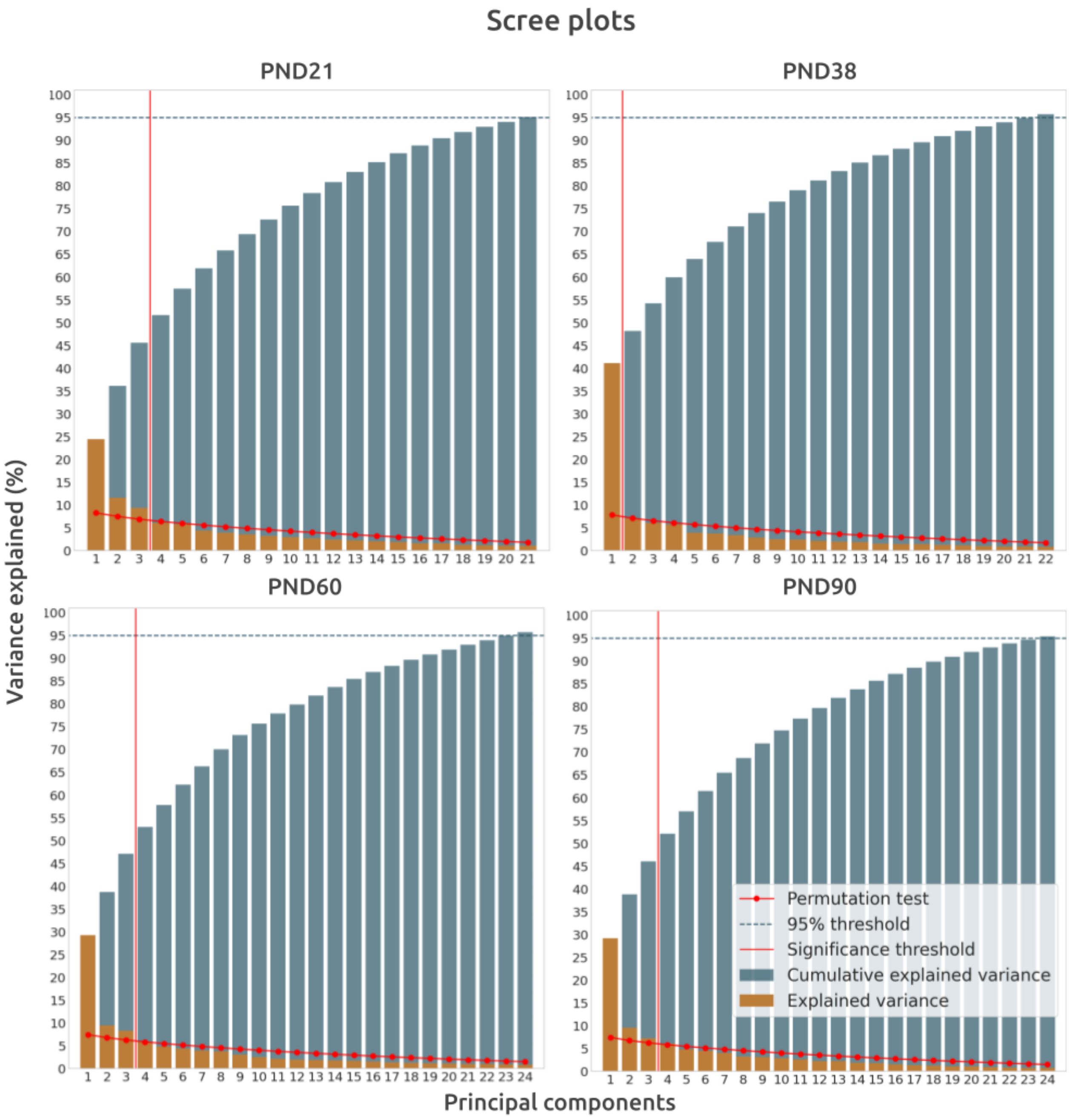


**Figure S1**: PCA scree plots. Variance explained for PCs up to 95% cumulative variance explained specific for individual timepoints.

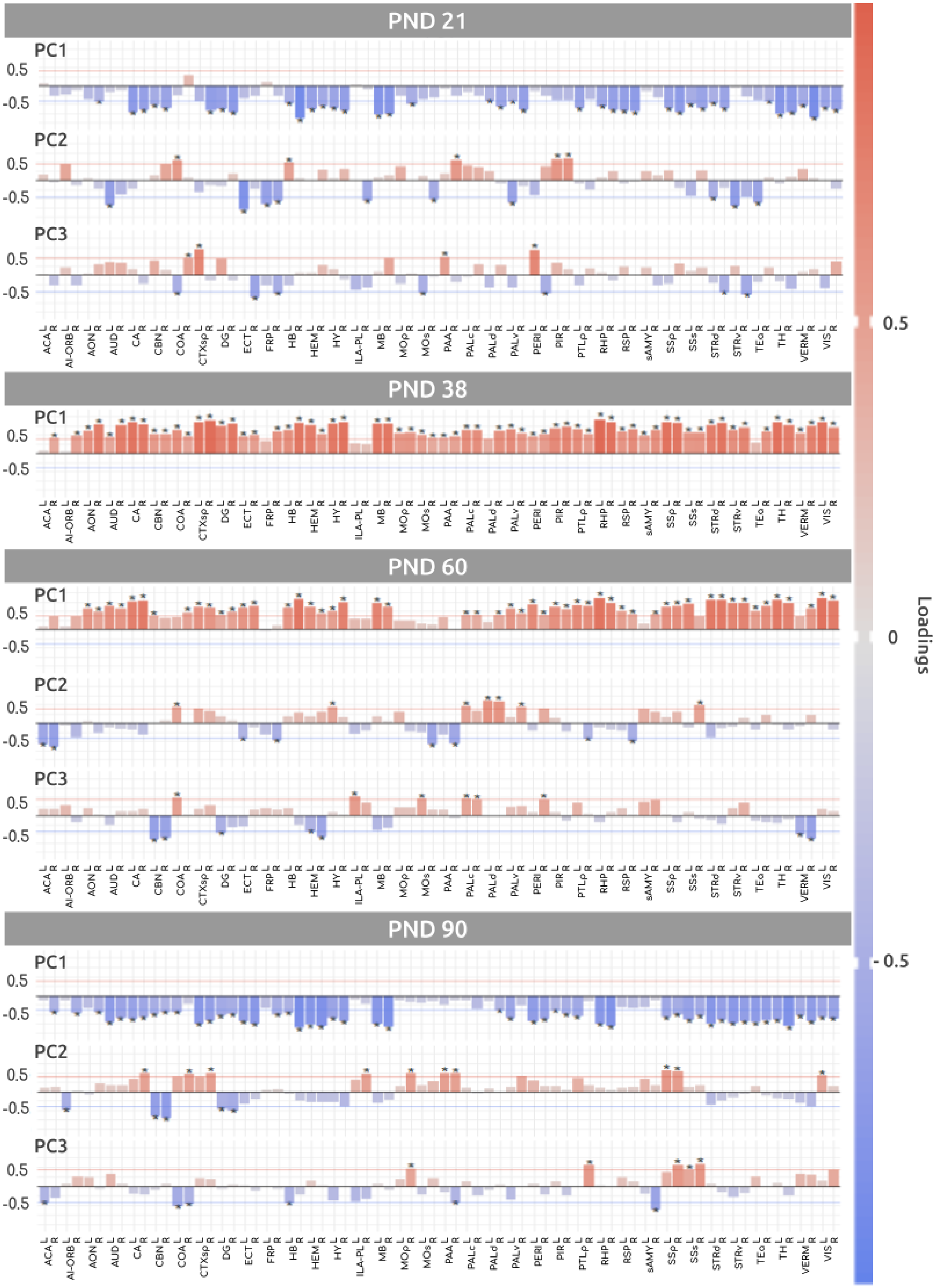


**Figure S2**: PCs brain regions loadings barplots. (* = Significant after permutation testing)

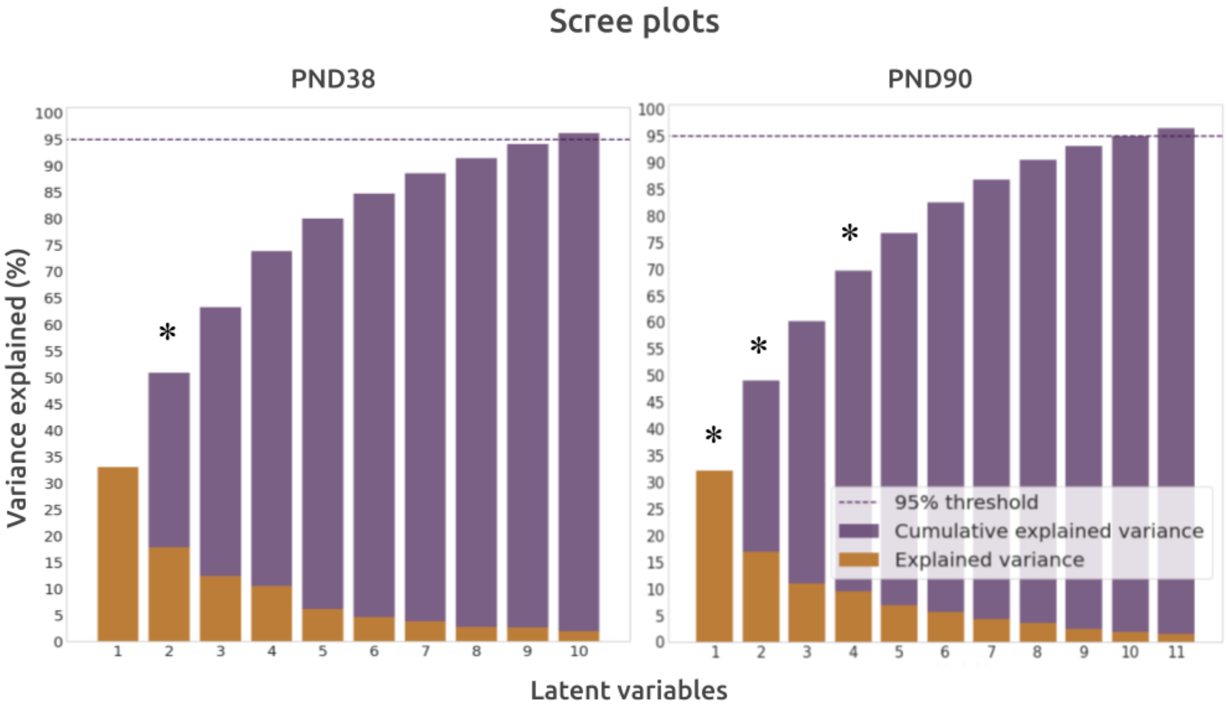


**Figure S3** : PLSC scree plots. (* = Significant after permutation testing)

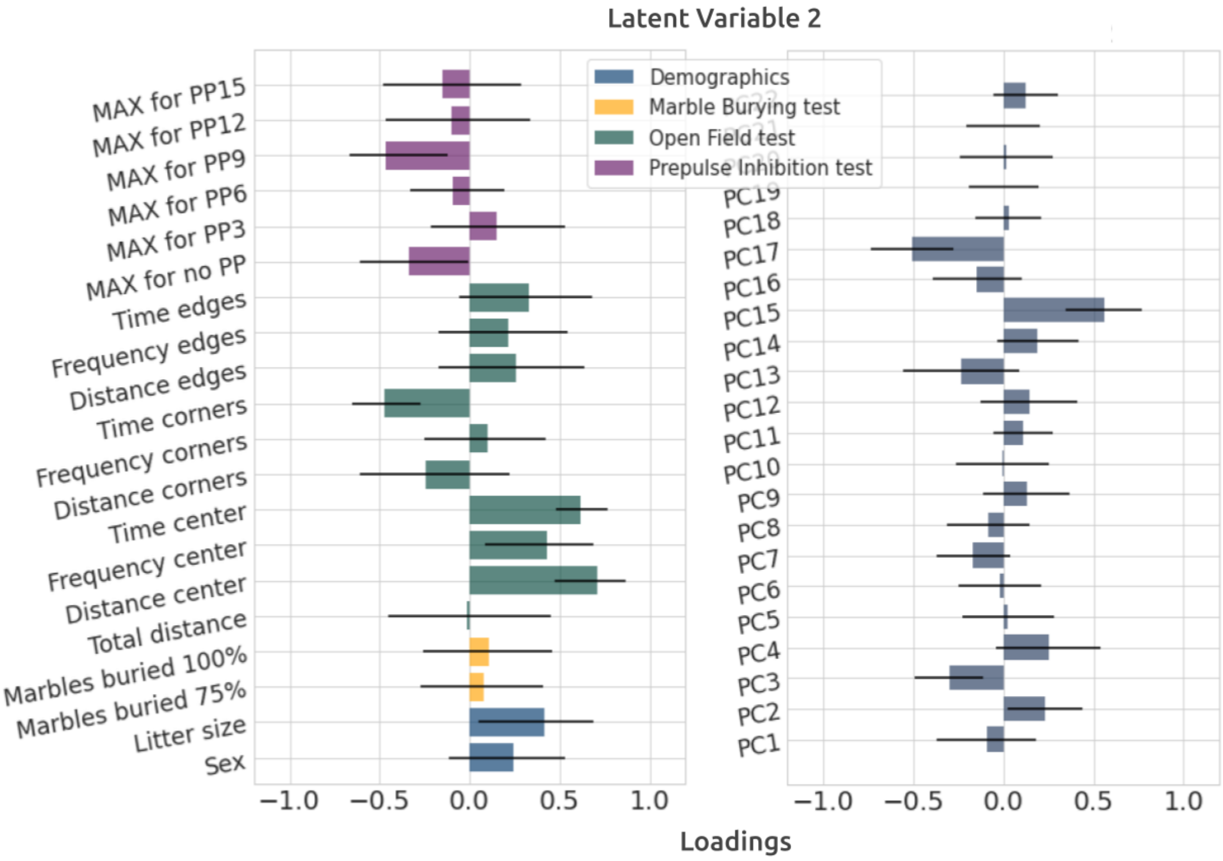


**Figure S4**: PND 38 latent variable 2 (LV2) behaviors and PC scores loadings.

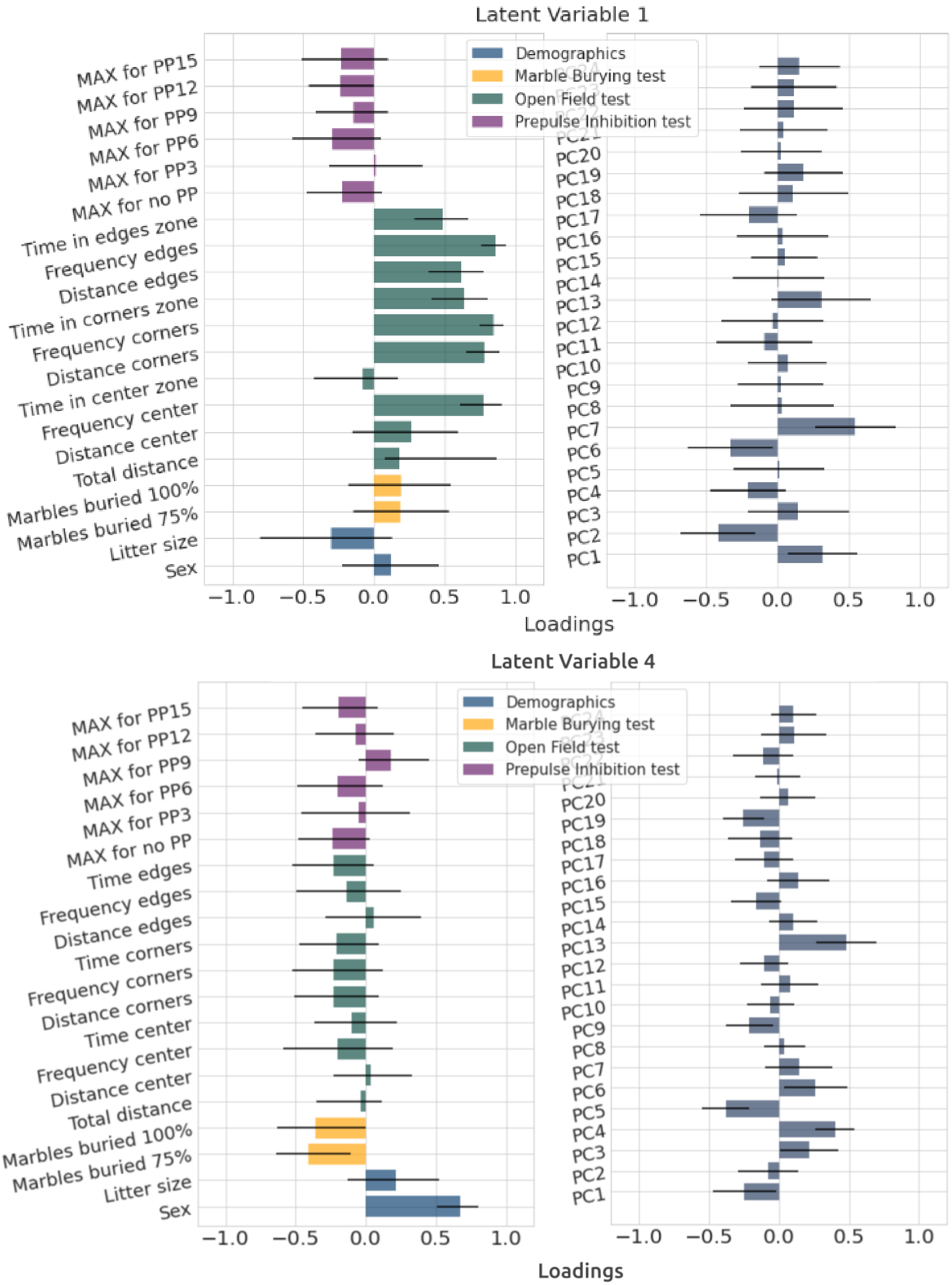


**Figure S5**: PND90 Latent variable 1 (LV1) and 4 (LV4) behaviors and PC scores loadings.

**Table S6:** Brain regions SS results overlap for model 3 (Age effect) and model 5 (Treatment*Age effect) power simulations respectively. These highlights the replication of an optimized sample size (smaller) when a greater number-of-litters of smaller size are selected for brain regions from different effect size ([0.20, 0.25], [0.40, 0.45]) simulations.

**Brain regions SS overlap between power simulations**

| **Age effect** | | | | | **Treatment * Age effect** | | | |
|---|---|---|---|---|---|---|---|---|
| **L** | | **R** | | **Regions** | **L** | | **R** | |
| **0.20** | **0.25** | **0.20** | **0.25** | | **0.40** | **0.45** | **0.40** | **0.45** |
| x | x | | x | ACA | x | x | x | x |
| x | x | | x | AI-ORB | | x | x | |
| x | x | x | x | AON | | x | | x |
| x | | x | x | AUD | x | | | |
| | | | x | CA | | | | |
| | | | x | CBN | | x | x | |
| x | x | x | | COA | x | x | | |
| | | | x | CTXsp | | x | x | |
| | x | x | | DG | | | x | x |
| x | | | x | ECT | | x | | |
| x | | | x | FRP | x | | | x |
| x | | | | HB | x | | x | |
| x | x | x | x | HEM | | | | x |
| x | | x | x | HY | x | | x | x |
| | | x | | ILA-PL | x | | | |
| x | x | | x | MB | | | | |
| x | x | | | MOp | | x | | x |
| x | x | x | x | MOs | x | | | x |
| | | | x | PAA | | | x | |

| | | | | | | | | |
|---|---|---|---|---|---|---|---|---|
| | | x | x | PALc | x | | | |
| x | | x | x | PALd | x | x | x | x |
| x | x | x | x | PALv | x | | | x |
| x | x | x | x | PERI | | | | |
| x | | x | | PIR | | x | x | x |
| x | | | | PTLp | x | | x | x |
| x | | | | RHP | x | | x | |
| x | | x | | RSP | | | | x |
| | x | x | | sAMY | | x | | |
| x | | x | | SSp | | x | | x |
| x | | x | | SSs | x | | x | |
| x | | | | STRd | | x | | |
| | x | x | | STRv | | x | x | |
| x | | x | | TEa | | x | x | |
| x | | x | | TH | | x | x | x |
| x | x | x | | VERM | x | | x | x |
| | x | x | x | VIS | | | | x |
